# Experience-dependent reorganization drives development of a binocularly unified cortical representation of orientation

**DOI:** 10.1101/761395

**Authors:** David Whitney, Jeremy T. Chang, David Fitzpatrick

## Abstract

Across sensory areas, neural microcircuits consolidate diverse streams of information into unified, representations of the external world. In the carnivore visual cortex, where eye-specific inputs converge, it has been posited that a single, shared columnar representation of orientation develops independent of sensory experience. In this study*, in vivo* calcium imaging with columnar and cellular resolution reveals a strikingly different developmental process in ferret visual cortex, starting with an early developmental period in which contralateral, ipsilateral or binocular stimulation each yield distinct well-organized representations of orientation that are misaligned at the columnar and cellular scale. Experience-dependent processes drive the reorganization of these three representations towards a single binocularly-aligned representation resembling the early binocular representation through concerted shifts in the preferred orientation of individual neurons. Thus, contrary to previous findings, a unified binocular representation of orientation results from an experience-dependent process that aligns the activity patterns of three distinct neural representations.

## Introduction

A significant step towards mature sensory processing is the developmental emergence of comprehensive neural representations of an organism’s environment. To achieve this, neural circuits combine information from different sensory sources to generate unified representations of the external world (Duhamel et al., 1998; King and Hutchings, 1987; Knudsen and Brainard, 1991; Wallace and Stein, 1997). The binocular and monocular representations of stimulus orientation in visual cortex provide an excellent example of this integrative process, since segregated inputs from the two eyes first come together to yield a unified binocular representation of stimulus orientation, such that visual stimuli presented to either eye yield similar population responses (Bridge and Cumming, 2001; Sarnaik et al., 2014) (Bridge and Cumming, 2001; Crair et al., 1998; Wang et al., 2010).

In carnivores and primates, the traditional view of orientation preference development, based on intrinsic signal imaging studies that lack cellular resolution, is that a spatially clustered, binocularly unified, and stable representation innately develops before visual experience (Chapman et al., 1996; Crair et al., 1998; Godecke and Bonhoeffer, 1996; Godecke et al., 1997; Sengpiel et al., 1998; White et al., 2001). Consistent with these findings, early modular patterns of spontaneous activity in ferret visual cortex, predict the mature orientation representation suggesting that a common network-scale representation develops before sensory experience (Smith et al., 2018). In contrast, studies in mice, which lack a structured network-scale organization, highlight the necessity of visual experience to generate feature-specific connectivity (Ko et al., 2013; Ko et al., 2014) and monocularly matched cellular orientation preferences (Gu and Cang, 2016; Wang et al., 2010). Because of this discrepancy, the role of sensory experience in the development of orientation representations in visual cortex, as well as the relationship between cellular- and network-scale representations across development remains unclear.

In this study, we examined the development of the unified binocular representation of orientation at both the cellular- and network-scales using widefield and two-photon calcium imaging in ferret visual cortex. We demonstrate that orientation-selective responses are already spatially clustered into distributed, modular representations before patterned visual experience. Despite this surprising degree of functional maturity, we discovered that cortical networks generate three distinct spatially-clustered representations of orientation preference at eye-opening: a monocular representation for each eye, and a third representation revealed through binocular stimulation. Furthermore, we demonstrate that shared binocular experience is necessary to drive network-scale alignment of these orientation representations towards a single binocularly unified representation. The absence of binocular experience during this period not only prevents network-scale alignment, but degrades the intrinsic modular organization of orientation preference into a state resembling the salt-and-pepper functional architecture of rodents. Finally, we observed that substantial changes in orientation preference, evident at both cellular- and network-scales, is a critical part of a dynamic process that leads to a unified representation of orientation preference resembling the early binocular representation. We propose that the emergence of a binocularly unified orientation representation in visual cortex reflects the experience-driven alignment of three distinct representations of orientation which arise independent of sensory experience.

## Results

### Mismatched monocular orientation preference is reduced across development

To measure monocular orientation preferences in the ferret visual cortex, we utilized viral expression of GCaMP6s with widefield epifluorescence imaging (Figure 1A-B). In ferrets, orientation selective responses are first observed around eye-opening (∼P30) and reach mature levels within one week (Chapman and Stryker, 1993; Chapman et al., 1996; Clemens et al., 2012; White et al., 2001). We first investigated the similarity of trial-averaged activity patterns evoked by drifting square wave gratings presented to either eye in visually experienced animals (experienced, P36-42, Figure 1A,C). The same stimulus presented to either eye evoked similar population responses (Supplemental Figure 1A-C) and produced a binocularly unified representation of orientation preference (Figure 1D). We next examined orientation-specific responses in visually naive animals prior to natural eye-opening (naive, P27-31, Figure 1A). In visually naive animals, stimuli presented to each of the opened eyes evoked distinct patterns of trial-averaged population activity (Figure 1E-F, Supplemental Figure 1D-F, Pearson’s correlation: Naive r=0.31±0.09 vs Experienced r=0.79±0.03 (Mean±SEM), n=6 vs 7 animals, p<0.001, unpaired permutation test), resulting in larger differences in monocular preferred orientation compared to experienced animals (Figure 1G, Mean monocular mismatch: Naive 27.91±4.19° vs Experienced 11.04 ±1.57° (Mean±SEM), n=6 vs 7 animals, p<0.001, unpaired permutation test). We failed to account for the monocular mismatch in naive animals with any rotational transform of the widefield orientation preferences, suggesting that monocular mismatch is cortical in origin and not due to physical torsion of the eyes (Supplementary Figure 1G-I, minimum monocular mismatch after allowing rotation: Naive 25.19±4.55° vs Experienced 9.23±1.03° (Mean±SEM), n=6 vs 7 animals, p<0.001, unpaired permutation test).

**Figure 1:**
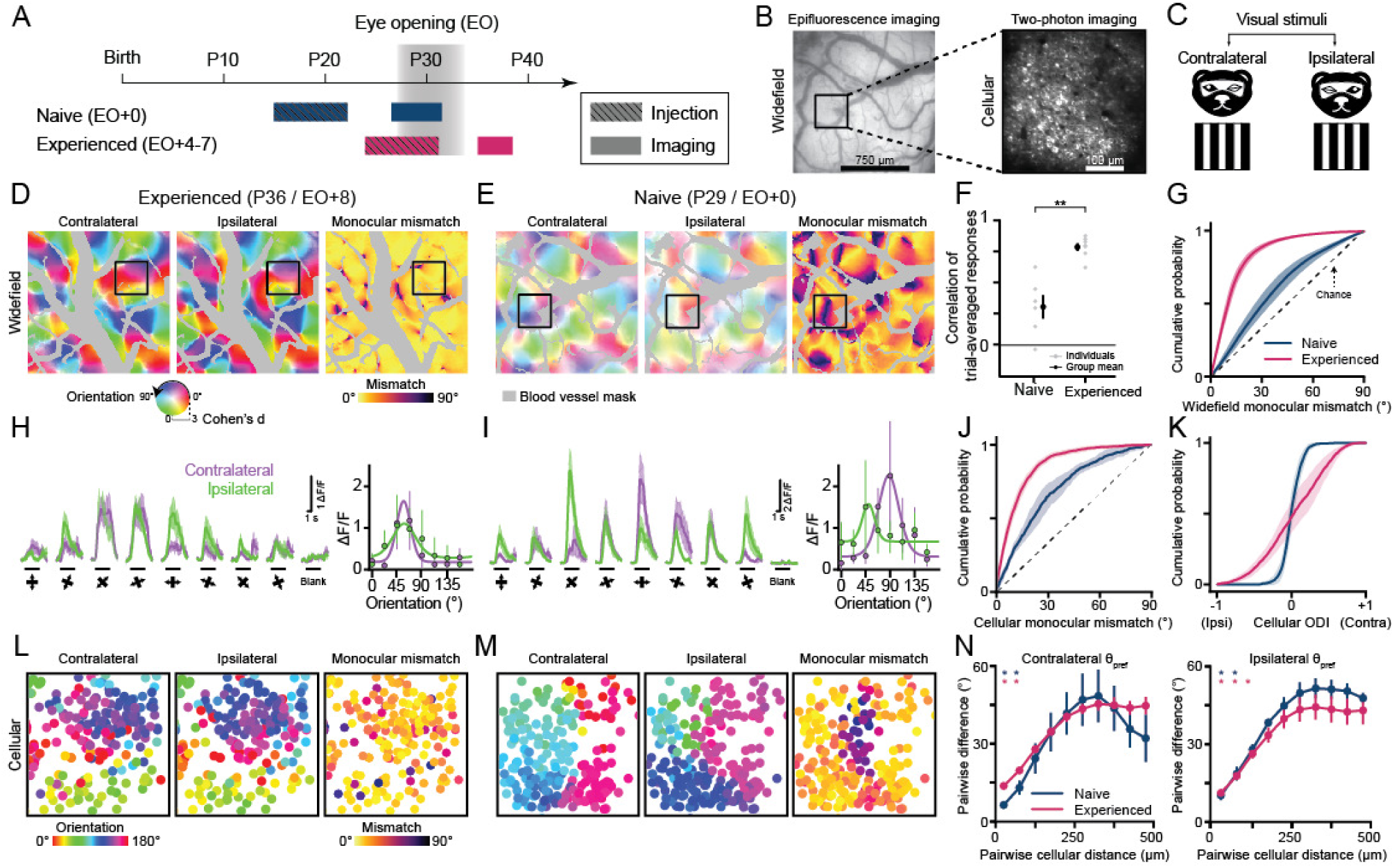
Mismatched monocular orientation preference is reduced across development. (A) Experimental timeline. (B) Widefield and two-photon images from the naive animal in (E). (C) Schematic of visual stimulation. (D-E) Widefield orientation preference maps for experienced (D) and naive animals (E). Far right column is monocular mismatch and grayed out regions represent blood vessels. Preferred orientation and selectivity (or monocular mismatch) are represented by image hue and saturation (or image color). (F) Pearson’s correlation between monocular trial-averaged responses. Gray and black dots reflect individual or grouped measurements. (G) Cumulative distributions of widefield monocular mismatch for naive and experienced animals (mean±SEM). (H-I) Cellular response traces (left panel, mean±SEM) and orientation tuning curves fit to trial-averaged responses (right panel, median±MAD) for experienced (H) and naive (I) animals. The black line under each trace indicates visual stimulation. (J-K) Cumulative distributions of cellular monocular mismatch (J) and ocular dominance index (K) for naive and experienced animals (mean±SEM). (L-M) Cellular orientation preference maps (and monocular mismatch maps) for the boxed regions in (D) and (E). Circles are cells and color represents the preferred orientation or monocular mismatch. (N) Distance-dependent, cellular clustering of the preferred orientation for naive and experienced animals (mean±SEM). Scale bars for widefield imaging and two-photon in (B) apply to (D-E, L-M). Asterisks denote significance between groups in (F) and distances where clustering is significant in (N) (*: p<0.05, **:p<0.01). Animal experiments: naive (n=3 cellular and n=6 widefield) and experienced (n=3 cellular and n=7 widefield).

Monocularly mismatched orientation preferences at the level of widefield epifluorescence imaging could arise within cortical networks in two possible scenarios. First, the ipsilateral and contralateral responses could reflect two distinct, but spatially intermingled neural populations, in which individual neurons respond to visual input from only one eye. Alternatively, neurons could be responsive to both eyes, but display different monocular orientation preferences. To distinguish between these possibilities, we used two-photon calcium imaging to measure responses to oriented stimuli in layer 2/3 pyramidal neurons of visual cortex in both naive and experienced animals (Figure 1A-B). We discovered that most recorded neurons were binocularly responsive (Figure 1H-I, Fraction of binocularly responsive cells: Naive 100±0% vs Experienced 96.27±1.57% (Mean±SEM), n=3 animals). However, unlike in experienced animals, the preferred orientation of binocularly responsive neurons in naïve animals often differed during monocular stimulation (Figure 1J, Supplementary Figure 2, Mean monocular mismatch: Naive 26.87±2.90° vs Experienced 13.11±1.03° (Mean±SEM), n=3 animals, p=0.002, unpaired bootstrap test). Furthermore, the relative response strength between the two eyes was comparable in binocularly responsive neurons of naïve animals suggesting that ocular dominance only becomes pronounced after eye-opening in experienced animals (Figure 1K, ODI: Naive 0.05±0.03 vs Experienced 0.23±0.06 (Mean±SD), Monocularity: Naive 0.09±0.04 vs Experienced 0.47±0.09 (Mean±SD), n=3 animals, p=0.04 for ODI and p=0.01 for Monocularity, unpaired bootstrap test). Taken together, our results demonstrate that neurons at eye-opening are binocularly responsive and frequently exhibit monocular mismatches in orientation preference that are rectified over the first week of visual experience.

To rule out the contribution of weaker orientation selectivity to the apparent orientation preference mismatch in naïve animals, we limited analysis to binocularly responsive neurons and pixels that exhibited well-tuned orientation responses (Fraction of well-tuned cells: Naive 56.76±3.94% vs Experienced 71.32±7.75% (Mean±SEM), n=3 animals; Fraction of well-tuned pixels: Naive 35.54±4.64% vs Experienced 80.87±2.33% (Mean±SEM), n=6 vs 7 animals). Furthermore, we evaluated whether the observed monocular mismatch could be accounted by measurement error. By bootstrap resampling the preferred orientation, we calculated the worst-case measurement error of the preferred orientation for either contralateral or ipsilateral stimulation. The observed monocular mismatch in naive animals was higher than the measurement error for either eye (Bootstrapped measurement error for cells: Naive contralateral 13.64° and ipsilateral 20.89°, Experienced contralateral 12.21° and ipsilateral 12.34°, n=3 animals; Bootstrapped measurement error for pixels: Naive contralateral 10.23±0.68° and ipsilateral 12.31±0.56°, Experienced contralateral 8.24±0.55° and ipsilateral 7.38±0.45° (Mean±SEM), n=6 or 7 animals). Therefore, weak orientation selectivity cannot explain the observed orientation preference mismatches.

In the mature visual cortex, neurons that share similar orientation preferences are spatially clustered, defining a modular, spatially distributed columnar representation of orientation (Figure 1D, Supplementary Figure 1A-C). The modular patterns evident in the trial-average population responses of naive animals visualized with widefield epifluorescence imaging (Supplementary Figure 1D-F) suggest that cellular orientation preferences are already spatially clustered (Figure 1L-M). To test this possibility, we quantified the spatial clustering by computing the cellular pairwise difference in preferred orientation as a function of distance. Naive animals did not display a higher degree of local scatter in preferred orientation than experienced animals for either eye (Figure 1N, p<0.05 within 100µm unpaired bootstrap test for both Naive and Experienced), consistent with the previously observed early columnar organization of spontaneous activity (Smith et al., 2018). Thus, we conclude that hallmarks of the mature visual cortex, such as orientation selectivity, binocular responsiveness, and a columnar architecture, are already present in visually naive ferrets. This high degree of functional network organization prior to eye-opening, makes the differences in monocular orientation preference even more surprising.

### Topologically distinct monocular orientation representations before eye-opening

We next sought to reconcile our observations that monocular orientation preferences appear to be well-organized spatially, but can be different for each eye. The presence of monocular mismatch within a cortical network displaying a fine-scale spatial clustering for the stimulus orientation of each eye raises the possibility that early binocular cortical networks harbor distinct modular representations of orientation for each eye, which are systematically misaligned in their preferred orientation (Figure 2A-C). To quantify whether the spatial layout of the preferred orientation is different for the monocular representations in naive animals, we computed the circular correlation of the monocular preferred orientations using widefield imaging. Consistent with our hypothesis, we found that the circular correlation of monocular preferred orientations was substantially lower in naive animals indicating that the monocular representations are distinct (Figure 2D, Circular correlation of monocular preferred orientations: Naive: r=0.44±0.11 vs Experienced: r=0.87±0.03 (Mean±SEM), n=6 vs 7 animals, p<0.001, unpaired permutation test). If the monocular mismatch does arise from distinct monocular representations, the spatial structure of the evoked response and the location of fine-scale topological features of monocular orientation preference maps, such as pinwheel centers and linear fractures, are unlikely to be shared. Because regions of low-homogeneity reflect pinwheels and linear fractures, we compared the homogeneity index (HI), which measures the local similarity of orientation preference within 100μm, for monocular orientation preference maps obtained with widefield imaging. Consistent with our hypothesis, we observed that the fine-network structures of the monocular representations, were less similar in naive animals than experienced animals (Figure 2C,E, Pearson’s correlation of monocular HI: Naive: r=0.41±0.11 vs Experienced: r=0.74±0.03 (Mean±SEM), n=6 vs 7 animals, p<0.001, unpaired permutation test).

**Figure 2:**
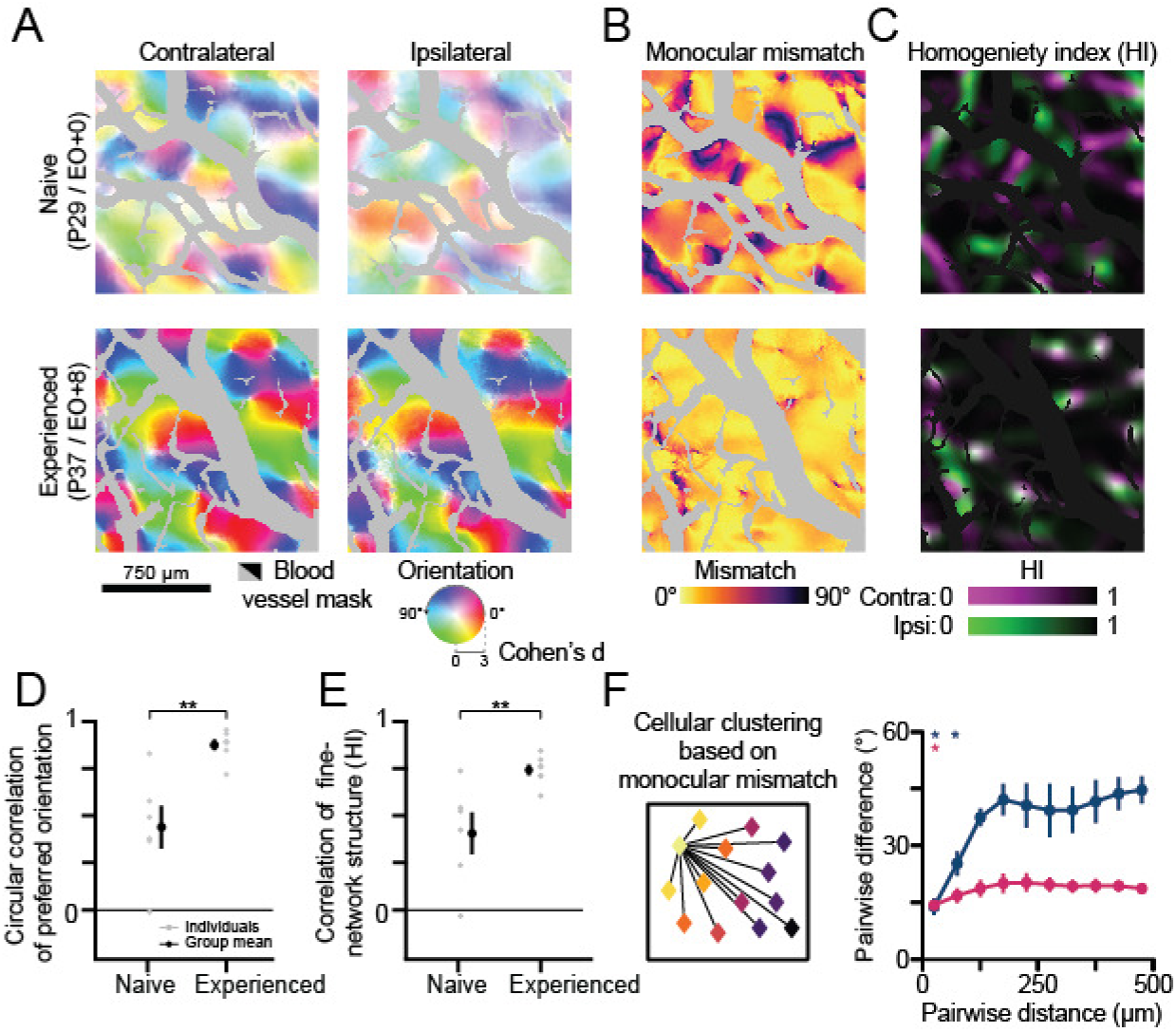
Monocular orientation representations are distinct before eye-opening. (A) Widefield orientation preference maps for naive (top) and experienced (bottom) animals. Preferred orientation and selectively are represented by image hue and saturation. Grayed or blacked out regions represent blood vessels. (B-C) Monocular mismatch (B) and homogeneity index (C) maps for the orientation preference maps in (A). (D-E) Circular correlation of preferred orientation (D) and Pearson’s correlation of fine-network structure (E, homogeneity index) for orientation preference maps in naive and experienced animals (mean±SEM). Gray and black dots reflect individual or grouped measurements. (F) Distance-dependent, cellular clustering of the monocular mismatch for naive and experienced animals (mean±SEM). Asterisks denote significance between groups in (D-E) and distances where clustering is significant in (F) (*: p<0.05, **:p<0.01). Animal experiments: naive (n=3 cellular and n=6 widefield) and experienced (n=3 cellular and n=7 widefield).

Clustering of differences is an expected consequence of the overlap of two well-organized representations. Therefore, another prediction consistent with two distinct monocular columnar representations, is that the degree of orientation preference mismatch would smoothly vary in a clustered manner across cortex. To test this hypothesis, we calculated the difference in observed monocular mismatch for pairs of cells as a function of distance. We observed that cells from naive animals displayed significant clustering of monocular mismatch in a region within 100μm (Figure 2F). We next considered the possibility that the clustering of monocular mismatch might be explained by the fact that our images are snapshots of an ongoing developmental process with aligned regions being more developmentally advanced than the misaligned ones. If so, we would expect to find a stronger negative correlation between a neuron’s orientation selectivity and monocular mismatch in naive animals compared to experienced animals. This is not what we observed (Pearson’s correlation between lowest monocular orientation selectivity versus monocular mismatch: Naive: r=-0.371±0.025 vs Experienced r=-0.317±0.017), indicating that the spatial clustering of monocular mismatch is not reflective of differences in the developmental stage of these cortical regions. Instead, regions of monocular mismatch appear to be a natural repercussion of equally mature misaligned columnar orientation representations. Together our findings suggest that innate factors before eye-opening establish distinct monocular representations of orientation, exhibiting unique topological structure, within the modularly, distributed cortical networks of visual cortex.

### Network-scale alignment of orientation representations requires patterned visual experience

Since the monocular preferred orientations are matched during the first week after eye-opening, we investigated whether this process requires patterned visual experience. We employed binocular eyelid suturing in a cohort of animals to delay eye-opening, depriving animals of patterned visual experience (deprived, P35-36, Figure 3A). Neurons from deprived animals demonstrated higher monocular mismatches in orientation preference than experienced animals (Figure 3B-D, Deprived 37.75±2.43° vs Experienced 13.11±1.03° (Mean±SEM), n=3 animals, p=.008, unpaired bootstrap test). Consistent with previous binocular deprivation studies (Chapman and Stryker, 1993; White et al., 2001), we also found that orientation selectivity in neurons from deprived animals was lower than from experienced animals (Figure 3E, p<0.05, unpaired bootstrap tests). Thus, patterned visual experience plays a dual role in the alignment of the monocular representations and the proper maturation of orientation selectivity.

**Figure 3:**
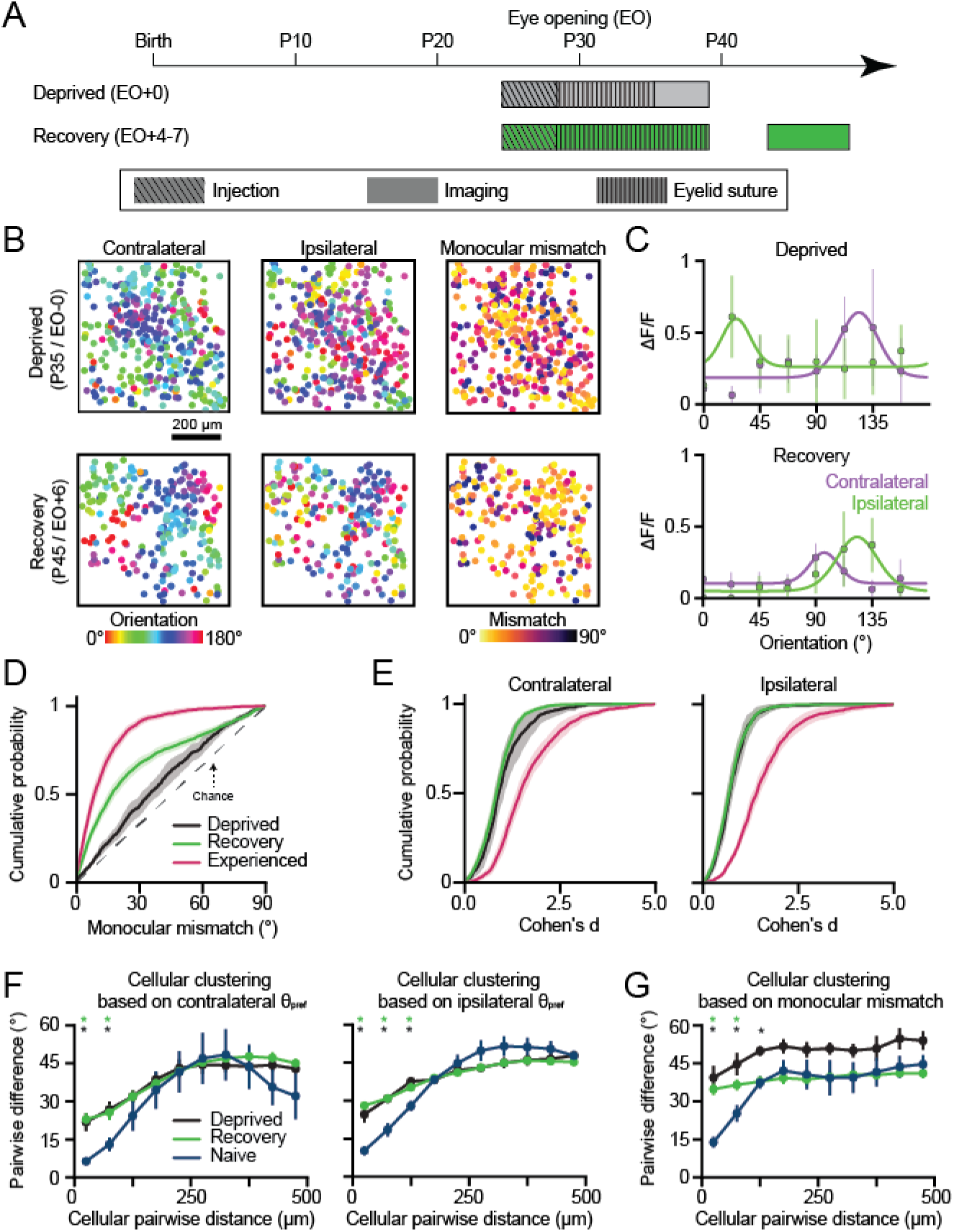
Network-scale alignment of orientation representations requires patterned visual experience. (A) Experimental timeline for deprivation and recovery. (B) Cellular orientation preference maps and monocular mismatch maps for deprived animals (top row) and recovery animals (bottom row). Circles are cells and color represents either the preferred orientation or monocular mismatch. (C) The trial-average response (median±MAD) and fit orientation tuning curves for cells from either a deprived animal (top row) or recovery animal (bottom row). (D) Cumulative distribution of cellular monocular mismatch for experienced, deprived, and recovery animals (mean±SEM). (E) Cumulative distributions of monocular orientation selectivity (Cohen’s d) for experienced, deprived, and recovery animals (mean±SEM). (F) Distance-dependent, cellular clustering of the preferred orientation for monocular stimulation in naive, deprived, and recovery animals (mean±SEM). (G) Distance-dependent, spatial clustering of monocular mismatch for naive, deprived, and recovery animals (mean±SEM). Asterisks in (F-G) denote distances where clustering is significant (*: p<0.05). Animal experiments: deprived (n=3), recovery (n=5), naive (n=3), and experienced (n=5).

Next, we investigated whether patterned visual experience during the normal first week of experience (P30-P37) is critical for producing a binocularly aligned orientation representation. In a cohort of animals, we delayed eye-opening by 9 days with binocular eyelid suturing, and then provided 4-7 days of patterned visual experience (recovery, P43-46, Figure 3A). Notably, recovery animals displayed deficits compared to normally reared animals, despite access to comparable amounts of patterned visual experience. Neurons from recovery animals displayed lower levels of orientation mismatch than those from deprived animals (Figure 3B-D, Recovery 27.00±1.43° vs Deprived 37.75±2.43° (Mean±SEM), n=5 vs 3 animals, p<0.05, unpaired bootstrap test), but orientation mismatches were significantly higher than from experienced animals (Figure 3D, Recovery 27.00±1.43° vs Experienced 13.11±1.03° (Mean±SEM), n=5 vs 3 animals, p<0.05, unpaired bootstrap test). Furthermore, cellular orientation selectivity in the recovery animals was indistinguishable from deprived animals (Figure 3E, p>0.05, unpaired bootstrap tests). Therefore, we conclude that visual experience, during the first week after natural eye-opening, plays an important role in the development of orientation selectivity and the alignment of monocular preferences, which cannot be recovered through comparable experience at a later developmental stage.

In addition to single-cell impairments, we discovered that the monocular representations of deprived and recovery animals showed systematically higher local scatter in preferred orientation than naive animals (Figure 3F-G, p<0.05 within 100µm unpaired bootstrap test vs Naive). Weakened spatial clustering implies that abnormal closed-eye experience after P30 induces a dramatic transformation of the representational network-structure in ferret visual cortex: from the modular functional organization characteristic of carnivores and primates into a more scattered functional organization reminiscent of the salt-and-pepper organization present in rodents. Alteration to circuit-level representations by abnormal experience is consistent with the underlying cortical circuitry becoming susceptible to visual experience at critical period onset (∼P30-P35)(Issa et al., 1999). Thus, the transition from closed-eye visual experience to patterned vision at ∼P30 is critically important for both the proper maintenance of spatially-clustered, columnar representations of orientation and the binocular alignment of preferred orientation.

### Binocular stimulation yields a third orientation representation before eye-opening

Sensitivity to visual experience raises the possibility that binocular experience, through simultaneous engagement of synaptic inputs from both eyes, plays a crucial role in establishing an aligned representation of orientation preference. Separate monocular representations of orientation preference could lead to binocularly-driven activity that is weaker or fundamentally different from monocularly-driven activity. Because of the potential importance of binocularly-driven activity and the observed monocular preference mismatch, we asked how the cellular-scale and network-scale responses evoked by binocular or monocular presentation of drifting gratings would compare (Figure 4A). We recorded responses to binocular and monocular stimulation using both two-photon (Figure 4B-E) and widefield (Figure 4F-G) calcium imaging in naive and experienced animals. As expected in experienced animals, the monocular and binocular cellular orientation preferences (Figure 4H-I, Mean orientation preference difference with binocular: Contralateral 10.44±1.10°, Ipsilateral 12.73±0.40° (Mean±SEM), n=3 animals) and network-scale orientation representations were well-matched (Figure 4J-K, Circular correlation with binocular: Contralateral r=0.93±0.01, Ipsilateral: r=0.93±0.02; Pearson’s correlation of homogeneity index with binocular: Contralateral: r=0.77±0.03, Ipsilateral: r=0.76±0.05 (Mean±SEM), n=7 animals).

**Figure 4:**
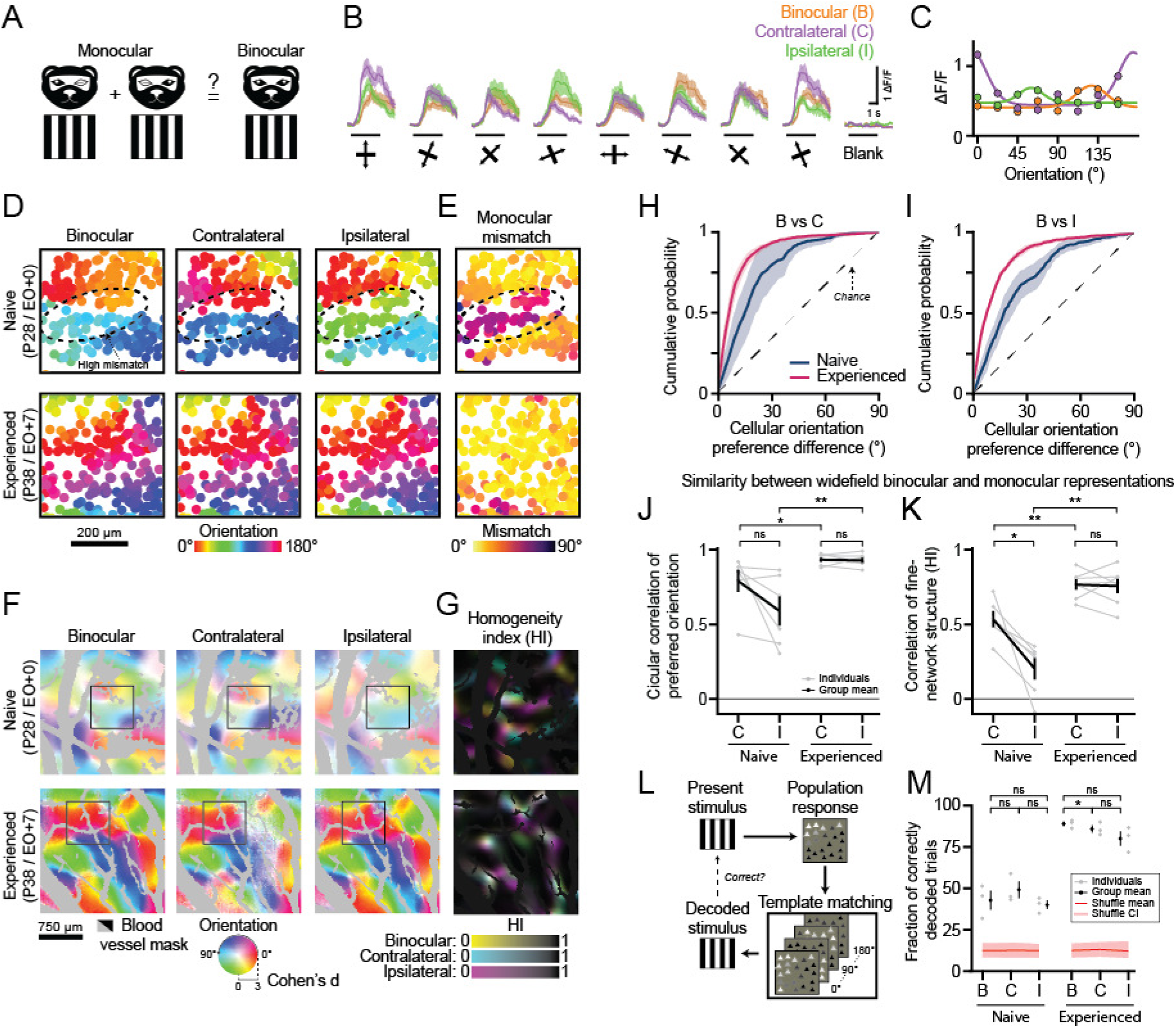
Binocular stimulation yields a third orientation representation before eye-opening. (A) Schematic of experimental paradigm. (B-C) Cellular response traces (B, mean±SEM) and orientation tuning curves fit to trial-averaged responses (C, median±MAD) in a naive animal. The black line under each trace indicates visual stimulation. (D-E) Cellular orientation preference (D) and monocular mismatch (E) maps for naive (top) and experienced (bottom) animals. Circles are cells and color represents the preferred orientation or monocular mismatch. (F-G) Widefield maps of orientation preference (F) and homogeneity index maps (G) for naive (top row) and experienced (bottom row) animals. Preferred orientation and selectively (or homogeneity index) are represented by image hue and saturation (or image color). Grayed or blacked out regions represent blood vessels, while boxed regions correspond to the cellular fields-of-view in (D-E). (H-I) Cumulative distributions of cellular orientation preference difference between the binocular and monocular preferred orientations (mean±SEM). (J-K) Circular correlation of the binocular to monocular orientation preference maps (J) and Pearson’s correlation of fine-network structure (homogeneity index) for the binocular to monocular preferred orientation maps (K) (mean±SEM). (L) Template-matching decoder schematic. (M) Fraction of correctly decoded trials to binocular and monocular stimulation (mean±SEM). Red line and shaded regions indicate the mean and 95% confidence intervals for shuffled data. Asterisks denote significance between groups (*:p<0.05,**:p<0.01, ns: p>0.05). Animal experiments: naive (n=3 cellular and n=6 widefield) and experienced (n=3 cellular and n=7 widefield).

In contrast, binocular stimulation in naive animals drove robust orientation selective responses (Figure 4B-C), even in neurons where the monocular mismatch was high, and the binocular and monocular preferred orientation frequently differed (Figure D-E, H-I). Systematic differences in the binocular and monocular preferred orientations in naive animals could result from the co-activation of differently tuned monocular inputs leading to a degraded binocular representation in regions of high monocular mismatch. Signatures of a degraded binocular representation would include reduced binocular orientation selectivity and reduced spatial clustering of preferred orientation. Spatial clustering of the binocular preferred orientation was comparable between naive and experienced animals (Supplementary Figure 3A, p<0.05 within 100µm for Naive and Experienced, unpaired bootstrap test), and smoothly mapped across the cortex for both the cellular-scale (Figure 4D) and network-scale representations (Figure 4E). Furthermore, at the cellular-scale, we failed to find any systematic difference between binocular and monocular orientation selectivity in either naive or experienced animals (Supplementary Figure 3B, p>0.05 for all comparisons, paired bootstrap test), and binocular orientation selectivity was only weakly correlated to the amount of monocular mismatch (Supplementary Figure 3C, Pearson’s correlation: Naive r=-0.25±0.12 vs Experienced r=-0.21±0.02 (Mean±SEM), n= 3 animals). Thus, we conclude that in naive animals cortical circuits generate a coherent representation of orientation preference in response to simultaneous binocular stimulation in spite of the mismatch in the monocular representations.

Next, we investigated whether early binocular responses reflect a distinct cortical representation or if they are more related to one of the monocular representations. At the cellular-scale, the binocular preferred orientation of naive animals was not biased to either of the monocular preferred orientations (Figure 4H-I, Cellular orientation preference difference with binocular: Contralateral 19.12±6.83°, Ipsilateral 23.40±4.29° (Mean±SEM), n=3 animals; Naive vs Experienced comparison: Binocular vs Contralateral p=0.104, Binocular vs Ipsilateral p=0.052, unpaired bootstrap test). We then tested whether the binocular preferred orientation could be predicted based on the strength of the monocular responses, and found that the binocular and monocular preferred orientation similarity was only weakly correlated to a cell’s ocular dominance (Supplementary Figure 3D, Pearson’s correlation with ODI in naive animals: Binocular vs Contralateral r=-0.20, p<0.001, Binocular vs Ipsilateral r=-0.20, p<0.001, n=3 animals). These results suggest that binocular stimulation gives rise to an orientation-specific response that is distinct from the constituent monocular responses.

Consistent with our cellular-scale findings, at the network-scale, preferred orientations of the binocular representation were not significantly more correlated with the preferred orientation of either eye in naive animals, but did trend towards being more similar to the contralateral eye representation (Figure 4J, Circular correlation of preferred orientation with binocular in naive animals: Contralateral r=0.79±0.05, Ipsilateral r=0.59±0.07 (Mean±SEM), n=6 animals, p=0.07, paired permutation test). The fine topological features of the binocular representation in naive animals more closely matched the contralateral representation suggesting that synaptic drive from the contralateral eye may preferentially influence binocular response (Figure 4K, Pearson’s correlation of HI with binocular in naive animals: Contralateral r=0.71±0.05, Ipsilateral r=0.46±0.07 (Mean±SEM), n=6, p=0.04, paired permutation test). Nonetheless, the preferred orientation and network structure of the binocular representation became significantly better matched to the monocular representations after patterned visual experience (Figure 4J-K, Naive vs Experienced comparisons: Circular correlation of preferred orientation with binocular: Contralateral: p=0.02, Ipsilateral: p<0.01; Pearson’s correlation of HI with binocular: Contralateral: p=0.03, Ipsilateral: p<0.01; n=6 vs 7 animals, unpaired permutation test). Together these results establish that the binocular representation of naive animals is not trivially related to either monocular representation and instead constitutes a third, distinct orientation representation.

Because three separate orientation representations emerge in the developing visual cortex prior to visual experience, we wondered if they were equally capable of encoding orientation-specific information at the cellular-scale. To probe this, we employed a template matching decoder to predict the stimulus orientation presented on each trial by comparing trial-evoked population activity patterns to the trial-averaged response pattern (Figure 4L)(Montijn et al., 2014). In naive animals, we could decode population responses from binocular and monocular stimulation at a rate higher than chance indicating that all three representations principally contribute to stimulus encoding (Figure 4M, Decoding performance in naive animals: Binocular 42.9±5.8%, Contralateral 49.5±5.0%, Ipsilateral 40.2±2.7% (Mean±SEM) vs chance 12.5%). Importantly, in naive animals, none of the orientation representations outperformed the others (p>0.05, paired bootstrap test), demonstrating that the binocular and monocular representations are comparable in decoding information about the stimulus orientation in naive animals.

### Shifts in cellular orientation preferences reduce monocular mismatch

So far, we have demonstrated that three distinct representations of orientation, two monocular and one binocular, emerge within the developing cortical circuitry before eye-opening, and that subsequent exposure to patterned visual experience drives functional reorganization towards a binocularly aligned representation. Visual experience could drive alignment in binocularly responsive cells by inducing shifts in the preferred orientation for one or more of the orientation representations. Alternatively, because we observed a continuum of mismatched cells, alignment could be achieved by driving the mismatched cells to become monocular, eliminating monocular mismatch and seeding the development of ocular dominance columns. To investigate which of these mechanisms contributes, in a cohort of animals we chronically tracked L2/3 neurons under two-photon imaging of GCaMP6s to monitor how the monocular and binocular orientation representations changed between eye-opening (naive) and four days after eye-opening (experienced) (Figure 5A-B, n=4 animals).

**Figure 5:**
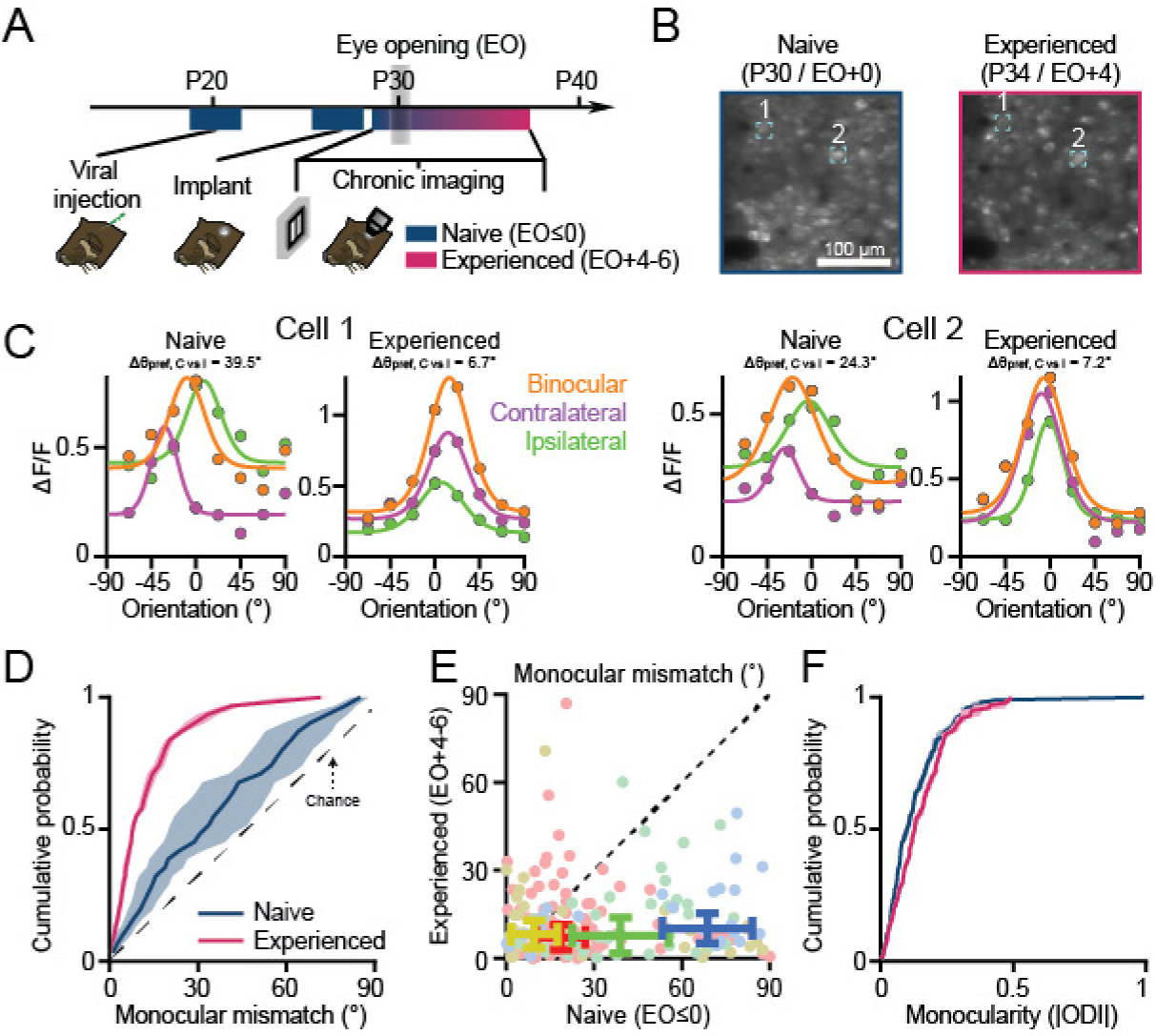
Shifts in cellular orientation preferences reduce monocular mismatch. (A) Experimental timeline for chronic imaging. (B) An example field-of-view of chronically tracked cells before and after eye-opening. (C) The trial-average response (median±MAD) and fit orientation tuning curves for chronically tracked cells. (D) Cumulative distributions of monocular mismatch for chronically tracked cells before and after eye-opening (mean±SEM). (E) Scatter plot of the monocular mismatch for chronically tracked cells before and after eye-opening. The error bars represent the Median±MAD for each animal experiment (n=4 animals), while the color of each dot or error bar represents the animal experiment. (F) Cumulative distributions of monocularity (or |ODI|) for chronically tracked cells before and after eye-opening (mean±SEM).

Roughly half of the chronically tracked neurons displayed well-fit orientation tuned responses to binocular and monocular stimulation at eye-opening (39.13±3.29%, Mean±SEM). Neurons that maintained orientation selective responsiveness across imaging sessions (73.53±7.97%, Mean±SEM) displayed systematic reductions in preference mismatch across all three representations (Figure 5C-F, Supplementary Figure 4, Mean monocular mismatch: Naive 35.47±6.09° vs Experienced 12.73±0.94° (Mean±SEM), p<0.001, paired bootstrap test) suggesting that shifts in preferred orientation at the cellular-scale contribute to orientation alignment. Furthermore, the monocularity of binocularly responsive neurons only modestly increased after four days of visual experience (Figure 5G, Monocularity: Naive 0.136±0.009 vs Experienced 0.154±0.003 (Mean±SEM), p<0.05, paired bootstrap test), including for the fraction of neurons that failed to maintain well-fit orientation selective responses after eye-opening (Monocularity: Naive 0.140±0.033 vs Experienced 0.247±0.09 (Mean±SEM), p=0.38, paired bootstrap test), and the degree of monocular mismatch at eye-opening did not predict the monocularity of individual neurons recorded days later (Bootstrapped Pearson’s correlation: actual r=0.03 [-0.12, 0.17] vs surrogate r=0.14 [-0.02, 0.30], Median [CI]). These results suggest that orientation alignment is driven primarily by systematic reorganization of the preferred orientation and that neurons likely become more monocular after the preferred orientation is matched (as previously shown in Figure 1H-I).

### Early orientation representations reorganize to a unified representation resembling the early binocular representation

We have demonstrated that there are three distinct representations of orientation at eye-opening that reorganize into a single coherent representation through changes in the orientation preference of individual neurons. In principle, the mature coherent representation could reflect a radical reorganization that leaves little trace of the representations that are present at eye-opening. To probe the nature of the reorganization, we asked whether the cellular-scale activity patterns at eye-opening exhibit any similarity to the binocularly unified representation that becomes evident days later. Our approach was to test whether a template matching decoder based on the trial-averaged response of the binocularly unified representation could be used to decode the presented stimulus orientation using trial-evoked population responses recorded at eye-opening. Under these conditions we found the decoder performed at rates that were above chance levels (Supplementary Figure 5A-C, Fraction of correctly decoded trials: Binocular 21.5±3.1%, Contralateral 17.92±2.0%, Ipsilateral 20.82±2.4% (Mean±SEM) vs Shuffles 12.5% [8.13, 16.88], Mean [CI]), although the decoding accuracy based on the experienced representation was substantially worse than the trial-average responses at eye-opening (p<0.001 for all comparisons, paired bootstrap test). Nevertheless, these results suggest that at the cellular-scale, experienced animals’ binocularly aligned representations bear a coarse resemblance to the representations evident at eye-opening.

To further probe this reorganization, we wondered whether there were any differences in the early representations that might provide insights into a possible dominance of one of the representations in shaping the later unified representation. From the 2-photon imaging experiments with a limited field of view, we noticed that at eye-opening, the range of preferred orientations found with binocular stimulation was greater than for monocular stimulation (Supplementary Figure 5H, Normalized difference from a uniform distribution: Binocular 0.51±0.05, Contralateral 0.65±0.02, Ipsilateral 0.66±0.03 (Mean±SEM); Binocular vs Contralateral: p=0.03, Binocular vs Ipsilateral: p=0.001, Contralateral vs Ipsilateral: p>0.05). These results raised the possibility that the size of orientation hypercolumns (the cortical distance required to represent the full range of orientations) is smaller for the binocular representation than the monocular representations at eye-opening.

To investigate this possibility further, we turned to chronic widefield imaging (n=12 animals, Supplementary Figure 6A), confirmed that the three representations were aligned following experience, and then used an established wavelet method to measure the width of orientation hypercolumns chronically (Supplementary Figure 6C,E-H)(Kaschube et al., 2010). Consistent with the single cell imaging, orientation hypercolumns at eye-opening were smaller for the binocular representation than the monocular representations (Supplementary Figure 6D, Orientation hypercolumn wavelength for Naive: Binocular 876.2±45.4μm, Contralateral 1021.7±54.2μm, Ipsilateral 1097.9±65.1μm (Mean±SEM); Binocular vs Contralateral: p<0.001, Binocular vs Ipsilateral: p=0.02, Contralateral vs Ipsilateral: p>0.05, paired bootstrap test). We then asked how hypercolumn size at eye-opening, compared with the size after the representations become aligned. By accounting for brain-related growth using landmark-based image registration, we found that orientation hypercolumns were overall smaller after eye-opening (Supplementary Figure 6D, Orientation hypercolumn wavelength for Experienced: Binocular 767.9±27.2μm, Contralateral 815.2±37.4μm, Ipsilateral 861.4±39.3μm (Mean±SEM); Naive vs Experienced: p<0.001 for all comparisons, paired bootstrap test), but the change in size of the hypercolumns for the binocular representation (∼12%) was almost half that of the monocular representations (∼20%).

Smaller changes in orientation hypercolumn width for the binocular representation suggest that the binocular representation may be more stable than the monocular representations (Figure 6C). Consistent with this hypothesis, we found that the layout of the preferred orientation for the binocular representation was significantly more stable than either of the monocular representations (Figure 6A,D, Circular correlation of the preferred orientation: Binocular r=0.40±0.04, Contralateral r=0.30±0.04, Ipsilateral r=0.16±0.04 (Mean±SEM), n=12 animals; Binocular vs Contralateral: p=0.02, Binocular vs Ipsilateral: p<0.01, Contralateral vs Ipsilateral: p=0.06, paired permutation test). If the binocular representation is more stable than the monocular representations, the fine-network structure of the binocular representation should remain more stable. Indeed, as assessed by HI, the fine-network structure of the binocular representation changes less than the monocular representations (Figure 6B,E, Pearson’s correlation of HI: Binocular r=0.40±0.06, Contralateral r=0.26±0.04, Ipsilateral r=0.12±0.07 (Mean±SEM); Binocular vs Contralateral: p=0.01, Binocular vs Ipsilateral: p<0.01, Contralateral vs Ipsilateral p=0.10, paired bootstrap test). These results demonstrate that the naive binocular representation is more stable than the monocular representations and suggests that after eye-opening the binocular unified representation might more closely resemble the naïve binocular representation.

**Figure 6:**
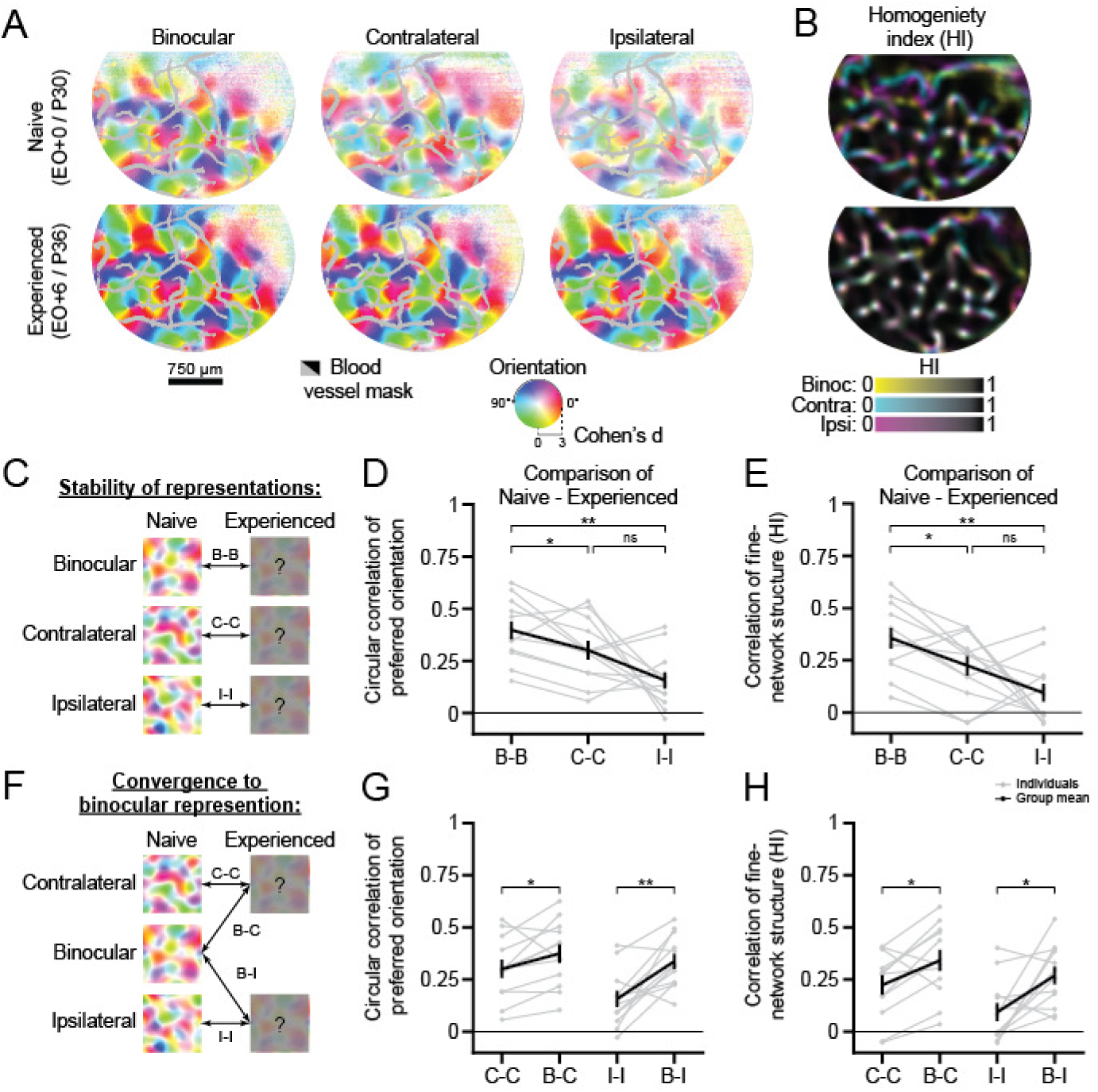
Early orientation representations reorganize to a unified representation resembling the early binocular representation. (A-B) Widefield maps of orientation preference (A) and homogeneity index maps (B) before and after eye-opening from a chronically imaged animal (top and bottom rows). Preferred orientation and orientation selectively (or homogeneity index) are represented by image hue and saturation (or image color). Grayed or blacked out regions represent blood vessels. (C) Schematic for determining the stability of the monocular and binocular representations before and after eye-opening. (D-E) Circular correlation of the preferred orientation (D) and Pearson’s correlation of fine-network structure (homogeneity index) (E) for comparing the stability of the binocular and monocular orientation preference maps before and after eye-opening (mean±SEM). (F) Schematic for determining if the experienced monocular orientation preference maps are more similar to the naive binocular or naive monocular orientation preference maps. (G-H) Circular correlation of the preferred orientation (G) and Pearson’s correlation of fine-network structure (homogeneity index) (H) for comparing the experienced monocular orientation preference maps either to the naive binocular or naive monocular orientation preference maps (mean±SEM). Gray and black lines/dots in (D-E, G-H) reflect either individual or grouped measurements (n=12 animals), while asterisks denote significance between groups (*:p<0.05,**:p<0.01, ns: p>0.05).

To directly test this hypothesis, we compared the experienced monocular representations to the naive binocular representation or corresponding naive monocular representation (Figure 6F). Indeed, we found that both the layout of preferred orientation and fine-network structure for the naive binocular representation was more similar to the experienced monocular representations (Figure 6G-H, Circular correlation of the preferred orientation: Experienced contralateral vs Naive contralateral r=0.30±0.04 or Naive binocular r=0.38±0.05, p=0.048, Experienced ipsilateral vs Naive ipsilateral r=0.16±0.04 or Naive binocular r=0.34±0.04, p<0.01; Pearson’s correlation of HI: Experienced contralateral vs Naive contralateral r=0.22±0.05 or Naive binocular r=0.34±0.05, p=0.02, Experienced ipsilateral vs Naive ipsilateral r=0.16±0.04 or Naive binocular r=0.34±0.05, p=0.02, paired bootstrap test). Taken together our results suggest that the naive binocular representation plays a dominant role in the experience-dependent reshaping orientation representations, likely reflecting the onset of coherent binocular experience that accompanies eye-opening.

## Discussion

### Three well-organized representations of orientation develop independent of experience

Our data demonstrates an unanticipated degree of functional organization in visual cortex before patterned visual experience. The prevailing view in mammals with a columnar architecture is that orientation selectivity is crudely established by eye-opening and that faint orientation preference maps are visible using intrinsic signal imaging (Chapman and Stryker, 1993; Chapman et al., 1996; Crair et al., 1998; White et al., 2001; Wiesel and Hubel, 1974). Fainter orientation preference maps visible at the network-scale could reflect the lower orientation selectivity of cells or an underdeveloped functional organization, such that well-tuned cells are not fully segregated into cortical columns. While orientation selectivity is weaker at eye-opening, we rule out an underdeveloped functional organization by demonstrating comparable spatial clustering of preferred orientation in naïve and experienced animals. In support of this view, chronic imaging in the developing ferret shows that the correlation structure of spontaneous activity patterns before eye-opening predicts the layout of the preferred orientation that becomes evident after eye-opening (Smith et al., 2018). Thus, our results reveal that well-organized columnar networks are present at eye-opening in ferret visual cortex, such that a fine-scale modular organization is evident across both cellular- and network-scales.

Yet surprisingly, these same well-organized networks generate different structured representations of orientation selectivity for monocular and binocular stimulation. These results challenge the prevailing view that at the onset of visual experience, segregated inputs from the eyes converge in visual cortex to yield a coarse, but binocularly matched representation of preferred orientation (Crair et al., 1998; Godecke and Bonhoeffer, 1996). Multiple factors may contribute to the different conclusions reached, such as species differences or reliance on intrinsic signal imaging, a less-sensitive technique for assessing preferred orientation. Regardless, no previous study has suggested that there could be an early developmental misalignment of organized columnar network structures. Indeed, across studies in carnivores, monocular and binocular stimulation have been historically used interchangeably, based on the assumption that monocular and binocular stimulation would produce comparable population responses in visual cortex (Chapman et al., 1996; Clemens et al., 2012; Krug et al., 2001; White et al., 2001). Our findings challenge this assumption, and highlight the complex interaction of eye-specific inputs, and the developmental shift in the similarity of monocular and binocular representations of orientation.

### Early binocular interactions are necessary for the development and maintenance of a single orientation representation

The presence of separate monocular and binocular representations of orientation selectivity before eye-opening suggests the developing cortex treats monocular inputs as separate streams of sensory information, and that shared binocular experience serves as an important catalyst for functional reorganization of the monocular representations towards a binocularly aligned representation. Our findings highlight the cooperative nature of ocular interactions in the generation of a binocularly unified orientation representation, which contrasts with most other studies that have focused on the competition of monocular inputs in the development of ocular dominance (Bienenstock et al., 1982; Blakemore and Van Sluyters, 1974; Katz and Shatz, 1996; Miller et al., 1989; Trachtenberg, 2015). Our deprivation experiments clearly show that patterned binocular visual experience is important for the alignment of the orientation representations, as well as the maintenance of spatially-clustered responses. This is consistent with growing evidence that binocular visual experience both during and after the critical period of visual development can improve visual acuity and correct deficits associated with amblyopia (Birch et al., 2015; Hess et al., 2010; Murphy et al., 2015), as well as computational studies suggesting that interocular correlations arising from natural binocular experience could play a crucial role in the alignment of the monocular orientation representations (Erwin and Miller, 1998; Miller and Erwin, 2001). Thus, shared sensory experience by the eyes appears crucial for the proper maturation of binocular interactions and the normal development of visual responses.

Importantly, binocular alignment of the monocular representations occurs during a critical time window when neural responses are still highly binocular. In the ferret, the critical period for ocular dominance begins about one week after eye-opening (Issa et al., 1999), suggesting that binocular alignment of orientation representations precedes the establishment of mature ocular dominance. Consistent with this hypothesis, we did not observe a dependence of the monocular matching process on the monocularity of cells. These results contrast with recent experiments in environmentally-enriched mice, where rescue of the monocular matching process in monocularly deprived animals depends on ocular dominance (Levine et al., 2017). Similarly, reverse-lid suture experiments in young kittens suggest that binocularly correlated visual input is unnecessary to develop matched monocular representations of orientation (Godecke and Bonhoeffer, 1996). Taken together these studies suggest that alignment of the preferred orientation in animals recovering from monocular deprivation, during a later developmental stage when cells are strongly monocular, benefit from the establishment of the dominant eye’s orientation representation. In contrast, over the course of normal development, monocular inputs cooperatively interact to achieve aligned orientation preference in an experience-dependent manner during the first week of visual experience.

### Experience-dependent reorganization across large-scale networks underlies alignment of orientation representation

Our experiments demonstrate that sensory experience plays a critical role in achieving a binocularly unified representation of orientation preference in primary visual cortex, challenging the conventional picture of a stable, experience-independent development of orientation preference in carnivores (Chapman et al., 1996; Godecke and Bonhoeffer, 1996; Godecke et al., 1997; Sengpiel et al., 1998). Instead, our data demonstrating experience-dependent development of a binocularly unified representation of orientation selectivity in ferret visual cortex bears more similarity to the binocular matching process reported in the developing mouse visual cortex (Gu and Cang, 2016; Wang et al., 2010; Yaeger et al., 2019). However, characterization in the mouse has been limited to single-cells, and has been unable to address whether the monocular mismatch reflects the misalignment of highly organized cortical representations. Given the salt-and-pepper organization of orientation preference in mouse visual cortex, it is also not clear how this level of characterization could even be evaluated. By leveraging a model species with a columnar architecture, we can now describe the binocular matching process of individual cells as reflecting the systematic alignment of network-scale representations, rather than a predictable byproduct of developing feature-specific synaptic connectivity from an initially non-specific organization (Ko et al., 2013; Ko et al., 2014).

Furthermore, the changes in the responses of individual neurons that are responsible for the binocular alignment of orientation preference have remained a mystery. To the best of our knowledge, this is the first study to chronically track the cellular changes underlying the normal alignment of sensory representations. We show that chronically tracked cells display shifts in preferred orientation reflecting reorganization of the orientation representations, rather than systematic loss of responsiveness to monocular input for one of the eyes in mismatched cells that could have accounted for the organization of ocular dominance. Across the neural population, our widefield epifluorescence imaging data demonstrates that the monocular representations undergo substantially more reorganization than the binocular representation. These results suggest that binocular vision after eye-opening instructs the development of a unified orientation representation. This conclusion contrasts with the prevailing views for alignment of sensory representations, which focus on either the experience-independent development of matched sensory representations or the experience-dependent alignment of one sensory representation to another (Cang and Feldheim, 2013; Cang et al., 2005; Chandrasekaran et al., 2005; Hyde and Knudsen, 2001; Knudsen and Knudsen, 1989; Smith and Trachtenberg, 2007; Triplett et al., 2009). Thus, our data suggests a novel solution to sensory representation alignment, where constructive interactions between segregated inputs lead to a third distinct representation, which serves as a template for the experience-dependent alignment of representations.

## Acknowledgements

We would like to thank J. Drayer and D. Ouimet for technical and surgical assistance, as well as members of the Fitzpatrick laboratory for helpful discussions. This research was supported by US National Institutes of Health grants EY011488 and EY026273 and the Max Planck Florida Institute for Neuroscience.

## Author contributions

All authors designed the study, analyzed the results, and wrote the paper. D.W. and J.C. performed the wide-field and 2-photon calcium imaging and contributed equally to this work.

## Methods

| REAGENT or RESOURCE | SOURCE | IDENTIFIER |
| --- | --- | --- |
| **Bacteria** |  |  |
| AAV2/1-hSyn-GCaMP6s-WPRE-SV40 | Chen et al., 2013 | RRID:Addgene_1100843 |
| **Experimental Models: Organisms/Strains** |  |  |
| Ferret | Marshall Farms | n/a |
| **Software and Algorithms** |  |  |
| MATLAB | MathWorks | https://www.mathworks.com/products/matlab.html |
| Python | Anaconda | https://www.anaconda.com/ |
| ImageJ | NIH | https://imagej.nih.gov/ij/ |
| Cell Magic Wand Tool | Wilson et al., 2017 | https://github.com/fitzlab/CellMagicWand |
| PsychoPy | Peirce, 2007 | http://www.psychopy.org |
| ScanImage | Pologruto et al., 2003 | http://scanimage.vidriotechnologies.com |
| Micromanager | Edelstein et al., 2010 | https://micro-manager.org/ |

### LEAD CONTACT AND MATERIALS AVAILABILITY

Further information and requests for resources and reagents should be directed to and will be fulfilled by the Lead Contact, David Fitzpatrick. This study did not generate new unique reagents.

### EXPERIMENTAL MODEL AND SUBJECT DETAILS

#### Animals

All experimental procedures were approved by the Max Planck Florida Institute for Neuroscience Institutional Animal Care and Use committee and were performed in accordance with guidelines from the U.S. National Institute of Health. Juvenile female ferrets from Marshal Farms were co-housed with jills on a 16-h light/8-h dark cycle.

### METHOD DETAILS

#### Viral injections and Eyelid Suturing

Viral injections were performed as previously described (Smith et al., 2018). Briefly we expressed GCaMP6s by microinjection of AAV2/1-hSyn-GCaMP6s-WPRE-SV40 (University of Pennsylvania Vector Core) into the visual cortex 6-14 days before imaging experiments. For eyelid suturing, eyelids were binocularly sutured during a short surgical procedure between P26-P30 using aseptic technique. Eyelid sutures were monitored daily until removed during an imaging experiment as noted in the main text.

#### Cranial window surgery

All animals were anesthetized and prepared for surgery as described above. In acute imaging experiments, skin and muscle overlying visual cortex were reflected and a custom-designed metal headplate (8mm DIA) was implanted over the injected region with MetaBond (Parkell Inc.). Then a craniotomy and a subsequent durotomy were performed, and the underlying brain stabilized with a custom-designed titanium metal cannula (4.5mm DIA, 1.5mm height) adhered to a thin 4mm coverslip (Warner Instruments) with optical glue (#71, Norland Products, Inc). The headplate was hermetically sealed with a stainless-steel retaining ring (5/16” internal retaining ring, McMaster-Carr) and glue (VetBond, 3M or Krazy Glue).

To allow repeated access to the same imaging field in chronic imaging experiments, cranial windows were implanted in each animal 2 days prior to the first imaging session. Using aseptic surgical technique, we adhered a metal headpost (7×7mm) to the skull ∼3.5mm anterior of bregma and a separate custom-designed, chamber implant overlying the injected region using MetaBond and black dental acrylic. At the end of the survival cranial window implant surgery, the metal cannula was sealed with a silicone polymer plug (Kwik-kast, World Precision Instruments) to protect the imaging window between imaging experiments. Whenever the imaging quality of the chronic cranial window was found to be suboptimal for imaging, the chamber was opened under aseptic conditions, any regrown tissue/neomembrane was removed, and a new coverslip was inserted.

For both acute and survival imaging experiments, eyelid sutures were removed or eyelids were separated where applicable to ensure visual stimulation was always presented to open eyes. Phenylephrine (1.25-5%) and tropicamide (0.5%) were applied to the eyes to retract the nictitating membrane and dilate the pupil, and the cornea was protected with regular application of eye drops (Systane Ultra, Alcon Laboratories). Prior to imaging, isoflurane levels were reduced from a surgical plane to ∼1-1.5%. After reaching a stable, anesthetic baseline for 30 minutes (∼280-300bpm), animals were paralyzed with pancuronium bromide (0.1 mg/kg/hr in lactated Ringer’s with 5% dextrose, delivered IV). Upon completion of imaging in acute experiments, isoflurane was raised to 5% and given Euthasol (0.5ml, IV). In survival experiments, animals were instead recovered from anesthesia and returned to their home cages. During recovery, up to 3 repeated doses of neostigmine (0.06mg/kg/hr, IV) and atropine (0.05mg/kg/hr, IV) were used to reverse paralysis.

#### Calcium Imaging Experiments

Widefield epifluoresence imaging of GCaMP6s was achieved with a Zyla 5.5 sCMOS camera (Andor) controlled by μManager (Edelstein et al., 2010). Images were acquired at 15Hz with 4×4 binning to yield 640×540 pixels through a 4x air-immersion objective (Olympus, UPlanFL 4x N/0.13NA). For analysis, images were spatially downsampled by a factor of 2× to yield 320×270 pixels at a spatial resolution of 11.63µm/pixel. Two-photon imaging of GCaMP6s was performed with a B-Scope microscope (ThorLabs) driven by a Mai-Tai DeepSee laser (Spectra Physics) at 940nm. The B-Scope microscope was controlled by ScanImage (Vidreo Technologies) in a resonant-galvo configuration with multi-plane images (512×512 pixels) being sequentially collected across either one or four imaging planes using a piezoelectric actuator for an effective frame rate of 30Hz or 6Hz respectively. Images were acquired at 2× zoom through a 16x water immersion objective (Nikon, LWD 16X W/0.80NA) yielding cellular fields-of-views of ∼0.7mm on each side (1.36µm/pixel).

#### Visual stimulation

Visual stimuli were delivered on a LCD screen placed approximately 25–30cm in front of the eyes using PsychoPy (Peirce, 2007). To evoke orientation responses, full-field square gratings at 100% contrast, at 0.015 or 0.06 cycles per degree and drifting at 1 or 4 Hz were presented at 8 or16 directions. In addition, “blank” stimuli of 0% contrast were also presented. Stimuli were randomly interleaved and presented for 3-4s followed by 3-6s of gray screen.

### QUANTIFICATION AND STATISTICAL ANALYSIS

#### General Analysis

ROI segmentation was performed manually in widefield epifluorescence imaging by manually drawing around cortical regions with robust visually evoked activity, while cellular ROIs were drawn using custom software in ImageJ (Cell Magic Wand)(Wilson et al., 2017). The baseline F_0_ for each pixel/cell was obtained by applying a rank-order filter to the raw fluorescence trace (25-30^th^ percentile) with a time window of 60s. The baseline corrected evoked activity was calculated as *(F-F_0_)/F_0_* = ΔF/F_0_. Grating evoked responses were then calculated as the difference of the average of the fluorescence ΔF/F_0_ over the full stimulus period (3-4s) and the average of the fluorescence ΔF/F_0_ over the pre-stimulus period (1.5-2s). Trial-averaged responses were calculated by taking the median across repeated trials to the same stimulus orientation.

#### Widefield Epifluorescence Imaging

For analysis of the wide-field epifluorescence imaging data, spatial filtering was necessary to eliminate signal strength variations and measurement noise. To determine an appropriate high-pass filter cutoff that did not fundamentally alter the underlying modular response, we first computed the average size (i.e. wavelength) of orientation hypercolumns for each imaging experiment using an established wavelet technique (Kaschube et al., 2010). We then spatially filtered each trial-evoked response image using a bandpass fermi-filter, where the low-pass filter cutoff was defined as 50µm and the high-pass filter cutoff was defined as the mean orientation hypercolumn wavelength across monocular and binocular stimulation. To maintain the original response amplitude of the population responses, spatially filtered images were ranged-fitted, such that the 5^th^ and 95^th^ percentiles intensity values matched the original, unfiltered images.

To evaluate orientation selective responses, we computed normalized single-condition response maps using the Cohen’s d metric. Pixels were considered well-tuned if they exhibited statistically significant orientation selectivity for both monocular and binocular stimulation using a permutation test. We excluded any pixels overlying blood-vessels that could be observed in the raw fluorescence images. Monocular mismatch was computed as the absolute angular difference between the contralateral and ipsilateral preferred orientations. To ensure that the measured monocular mismatch for the population could not be artifactually explained by high variation in the preferred orientation, we evaluated the worst-case measurement error. Here we bootstrap resampled trial-averaged responses using replacement and recomputed the preferred orientation (n=1,000). The measurement error for each pixel was defined as the median error of the bootstrapped preferred orientations with the measured preferred orientation. We then defined our worst case measurement error as the value of the pixel exhibiting large measurement error (95^th^ percentile).

Orientation preference differences between the monocular and binocular preferred orientations were computed using a similar formula to the monocular mismatch. The circular correlation was used to assess differences in the layout of preferred orientation for monocular and binocular orientation maps. For computing the similarity of the fine network structure in orientation preference maps, we computed the Homogeneity Index (Wilson et al., 2017). We then quantified the similarity between the Homogeneity Index maps for the monocular and binocular orientation maps using Pearson’s correlation.

##### Two-Photon Imaging

Cells were considered responsive if the trial-averaged response to any stimulus orientation for either monocular or binocular stimulation was greater than two standard deviations of the response to a blank stimulus. The preferred orientation for responsive cells was assessed by fitting a Von Mises function to the trial-averaged responses. Cells were considered well-tuned if the R^2^ value of the orientation fits to monocular and binocular stimulation were all ≥0.5. To evaluate a cell’s orientation selectivity, we used the Cohen’s d metric and compared the responses of the preferred and orthogonal orientations. Spatial clustering of the preferred orientation or monocular mismatch was assessed by calculating the pairwise correlation for cells located within a fixed range from each other, ranging from distances of 50 µm to 500 µm. To evaluate the degree that all preferred orientations are represented uniformly within each of our cellular fields, we estimated the normalized difference from a uniform distribution, where the values range from 0 (all orientations equally represented) to 1 (only a single representation represented). Monocular mismatch, orientation preference differences, measurement error, and the homogeneity index were computed using the same formulas previously described used for pixels (*see widefield imaging section*). To evaluate the degree that monocular input from one eye consistently drove stronger responses, we computed the ocular dominance index (ODI), where values range from −1 to +1, 0 indicates a lack of bias, and −1 and +1 values indicate complete dominance from either the ipsilateral or contralateral eyes. Monocularity was defined as the magnitude of the ODI, with values ranging from +0 (binocular) to +1 (monocular).

To decode the stimulus orientation from the population activity vectors recorded during each trial, we used a normalized template matching algorithm (Montijn et al., 2014). To ensure our analysis was comparable across different fields-of-view where the number of cells present varies widely, we limited the number of cells to 50 at any time for our decoder. The similarity metric computed for each template was bootstrap resampled without replacement, but a different subset of cells was used for each resampling procedure (n=1,000). The decoded stimulation orientation was defined as the stimulus orientation for the template showing the highest median similarity across bootstrap resampling to the population activity vector.

##### Statistics

Unless otherwise noted, hypothesis testing was performed using either a permutation or bootstrapping test (p<0.05). All statistical tests were two-tail, except when assessing if the performance of the normalized template matching decoder was above chance-levels (one-tail). Unequal group sizes were maintained for unpaired tests, while resampling was restricted between paired measurements for any paired test.

## DATA AND CODE AVAILABILITY

Data and code used in this study are available upon reasonable request from the lead contact.

---

**Supplemental Figure 1:**
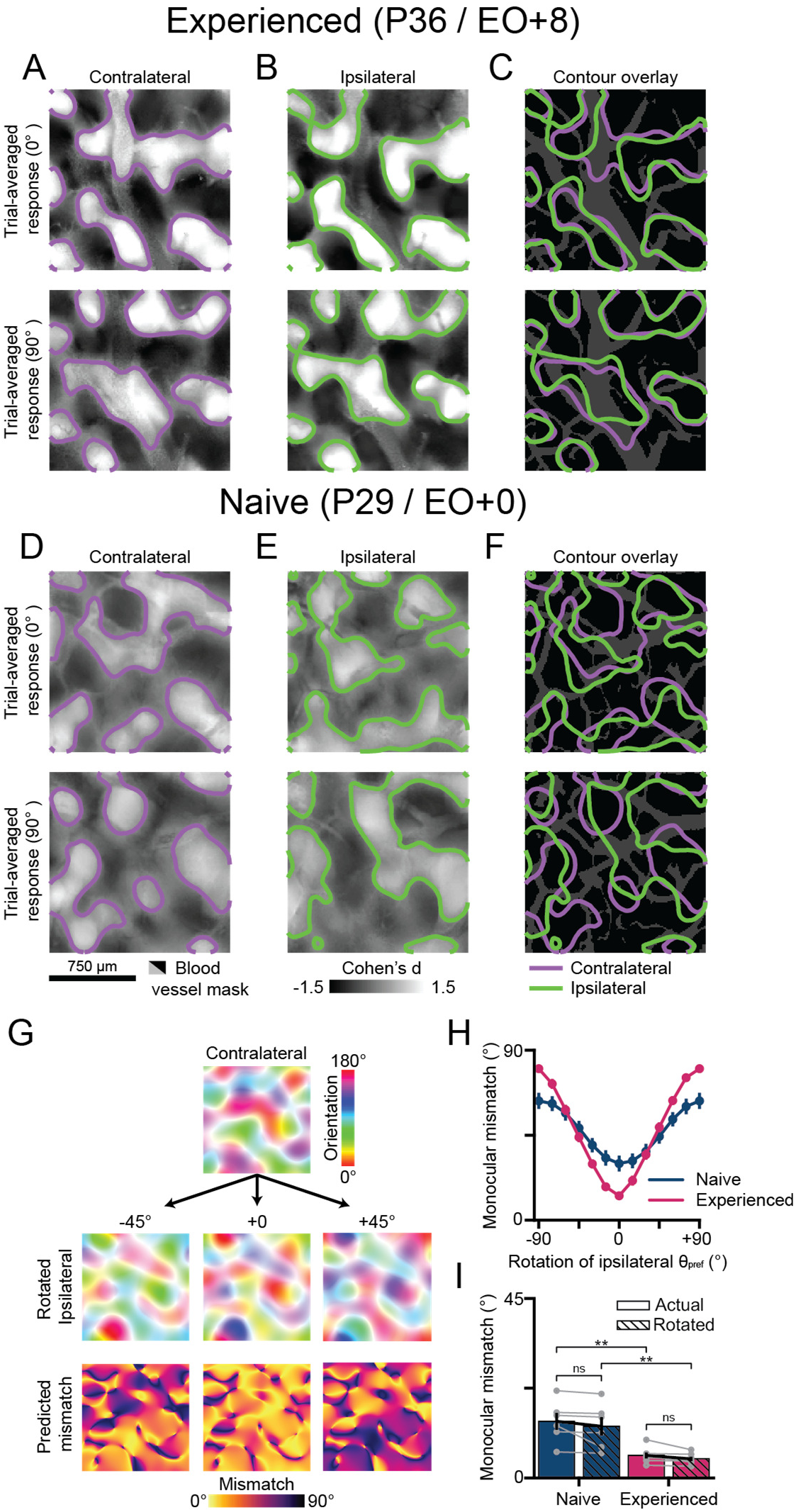
Activity patterns evoked by monocular visual stimuli yield different activity patterns in visually naive animals. (A-B) Trial-averaged, cocktail-blank subtracted response maps evoked by monocular presentation to drifting gratings for the experienced animal in Figure 1E. Contours denote active domains. Active domains were defined as regions where the relative activity of z-scored trial-average response maps was >0.25 after applying a 10×10 median filter. (C) Contour overlay of the active domains in (A-B). (D-F) Same as (A-C), but for the naive experienced animal in Figure 1D. For all widefield images, grayed or blacked out regions represent blood vessels. (G) Schematic of rotating the ipsilateral preferred orientation map relative to the contralateral preferred orientation mapn. (H) The effect that rotating the preferred orientation of the ipsilateral eye has on the monocular mismatch in widefield imaging across naive and experienced animals (mean±SEM). (I) The widefield monocular mismatch before and after rotating the ipsilateral orientation preference map to minimize the monocular mismatch. Gray and black dots respectively reflect individual or grouped measurements. Asterisks denote significance between groups (ns: p>0.05, **:p<0.01). Animal experiments: naive (n=6) and experienced (n=7).

**Supplemental Figure 2:**
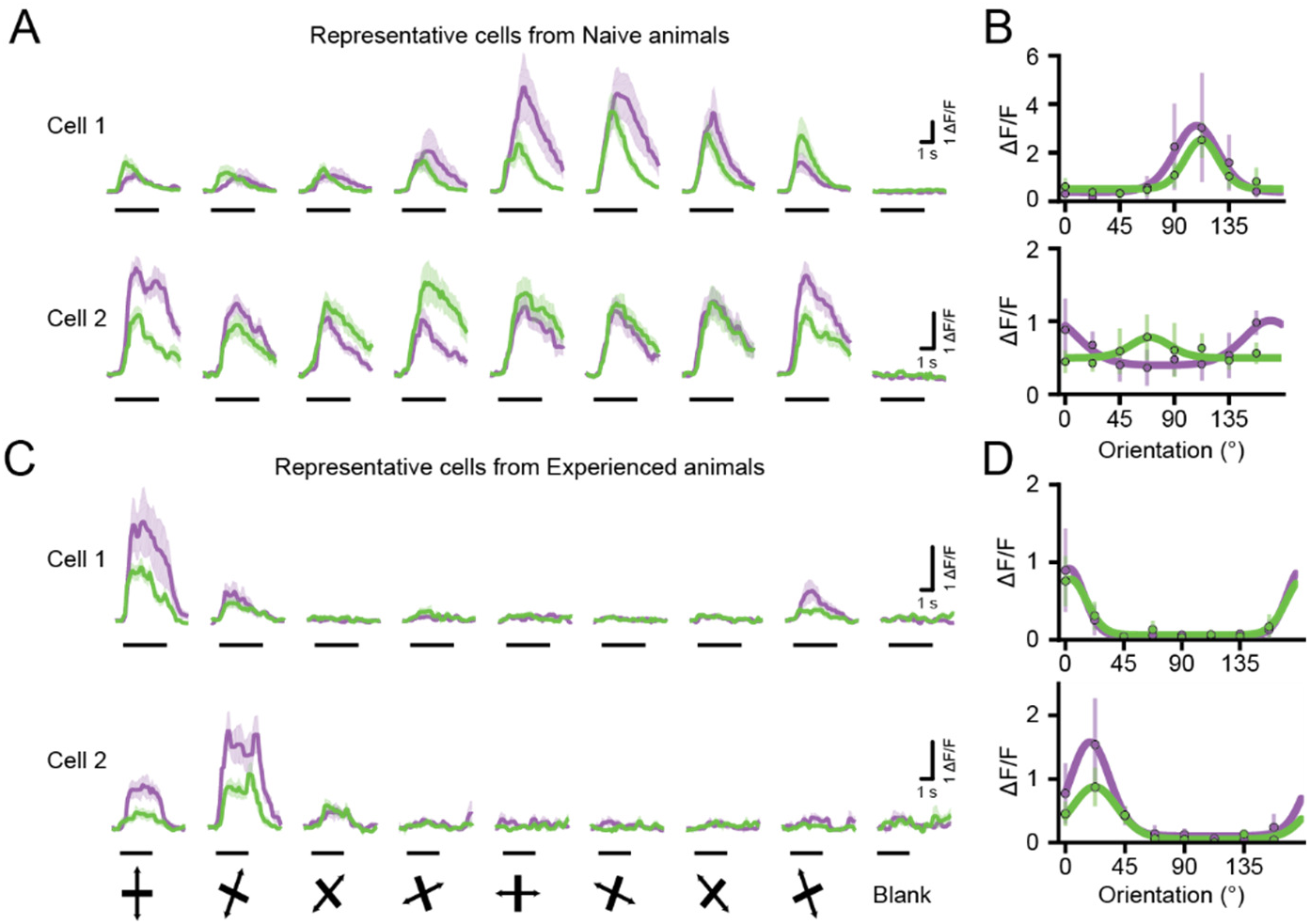
Additional examples of cellular responses in visually naive and experienced animals. (A) Representative cellular response traces from naive animals (mean±SEM). The line under each response trace indicates visual stimulation period. (B) For cells in (A), the trial-average responses (median±MAD) and fit orientation tuning curves. (C-D) Same as (A-B), but for cells from experienced animals.

**Supplemental Figure 3:**
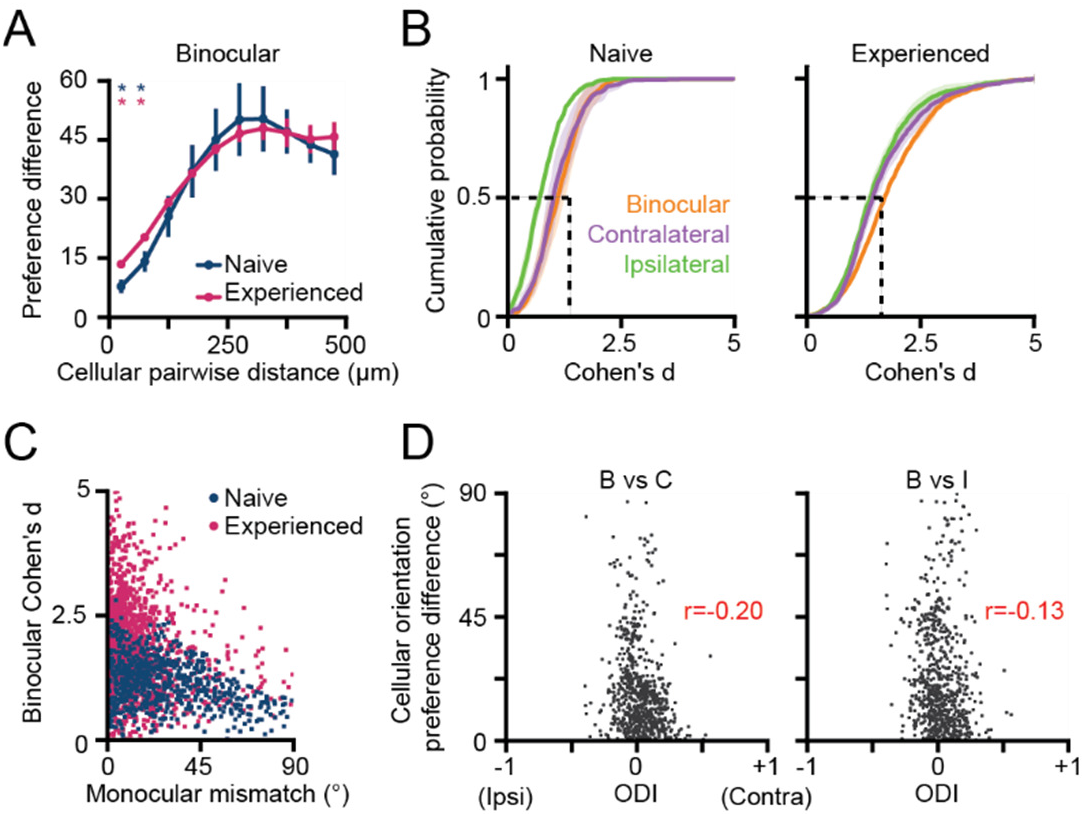
Additional analyses of the binocular and naive responses. (A) Distance-dependent, cellular clustering of the binocular preferred orientation for naive and experienced animals (mean±SEM). (B) Cumulative distributions of binocular and monocular orientation selectivity (Cohen’s d) for naive and experienced animals (mean±SEM). (C) Scatter plot of the cellular monocular mismatch and binocular selectivity (Cohen’s d) for naive and experienced animals. (D) Scatter plots of ODI and the orientation preference difference of the binocular preferred orientation with either monocular preferred orientation. The Pearson’s correlation coefficient (r) is listed in the figure panel. Animal experiments: naive (n=3 animals) and experienced (n=3 animals).

**Supplemental Figure 4:**
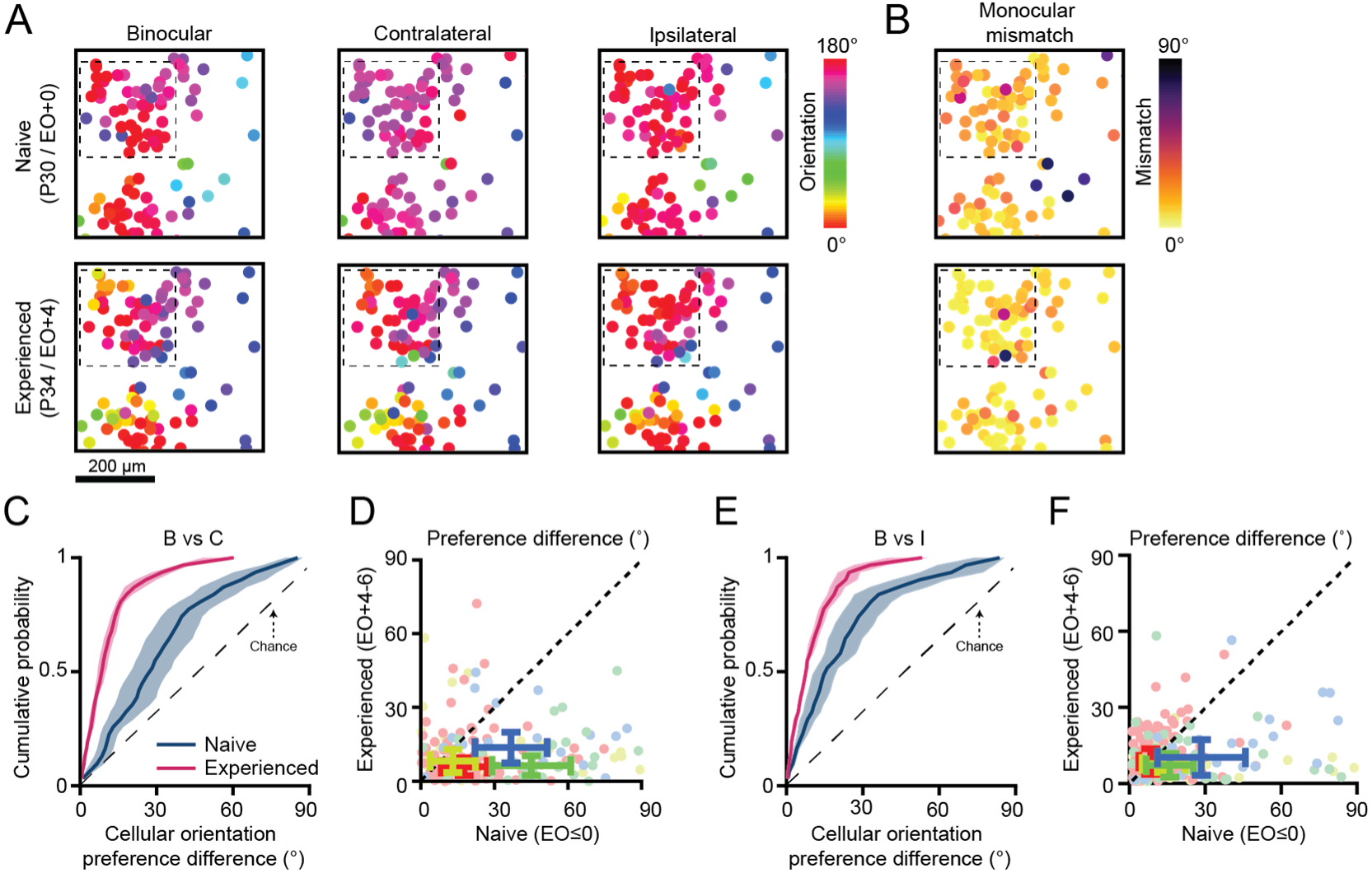
Additional quantification of binocular matching in chronically tracked cells. (A-B) Representative cellular orientation preference maps (A) and monocular mismatch maps (B) for chronically tracked cells before (top row) and after (bottom row) eye-opening. Circles are cells and color represents either the preferred orientation or monocular mismatch. (C) Cumulative distributions of orientation preference difference of the binocular and contralateral preferred orientations for chronically tracked cells before and after eye-opening (mean±SEM, n=4 animals). (D) Scatter plot of the orientation preference difference of the binocular and contralateral preferred orientations for chronically tracked cells before and after eye-opening. The error bars represent the Median±MAD for each animal experiment (n=4 animals), while the color of each dot or error bar represents the animal experiment. (E-F) Same as (C-D), but for the orientation preference difference of the binocular and ipsilateral preferred orientations.

**Supplemental Figure 5:**
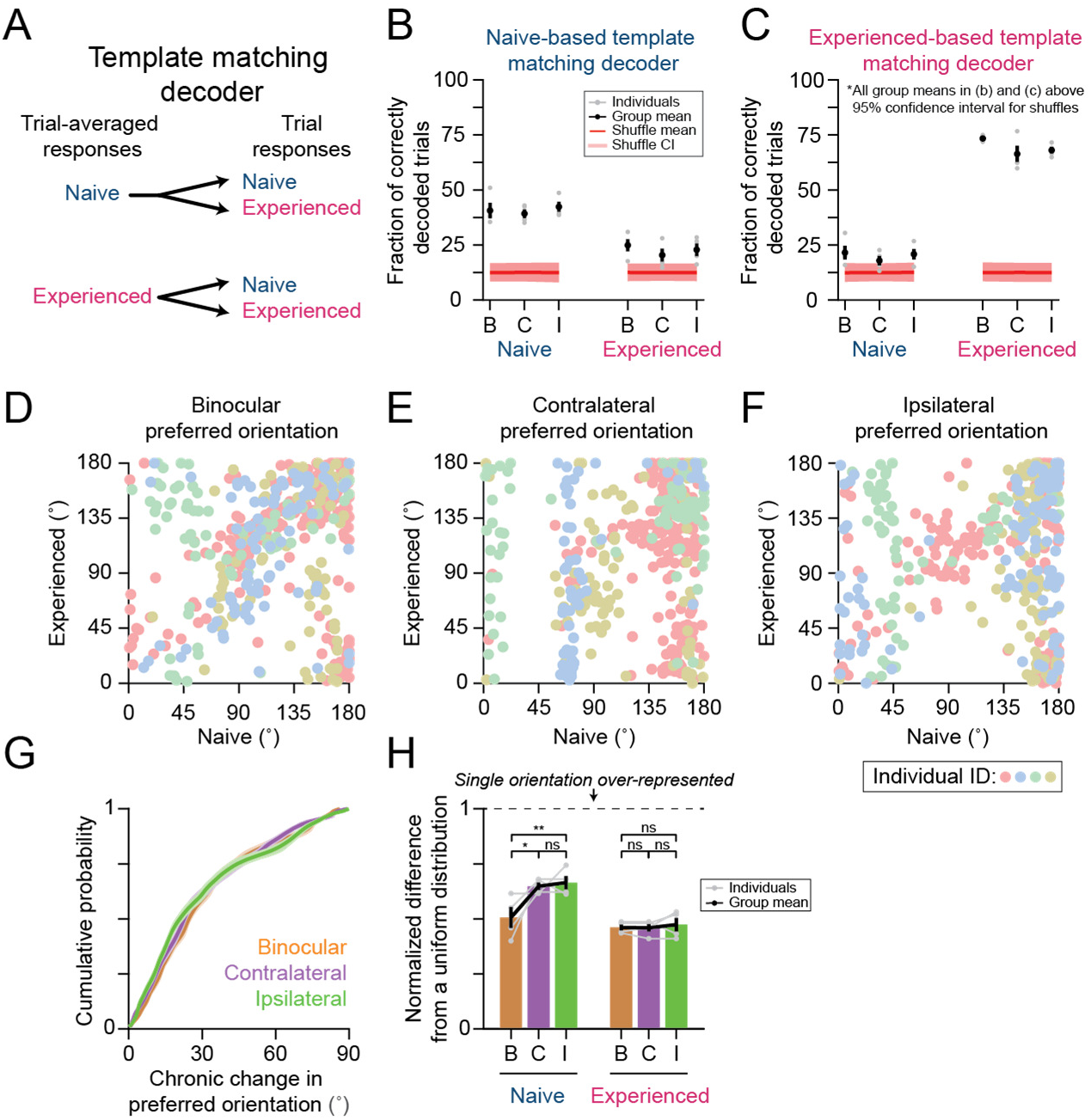
Additional quantification of chronically tracked cells. (A) Template-matching decoder schematic for chronically tracked cells. (B) Fraction of correctly decoded trials to binocular and monocular stimulation before and after eye-opening using decoder templates generated before eye-opening (mean±SEM). (C) Same as (B), but using decoder templates generated after eye-opening. (D-F) Scatter plot of the binocular (D), contralateral (E), and ipsilateral (F) preferred orientations for chronically tracked cells before and after eye-opening. (G) Cumulative distributions of orientation preference difference across time for chronically tracked cells before and after eye-opening (mean±SEM). (H) Normalized average difference of the distribution of preferred orientations in a cellular field-of-view from a uniform distribution of preferred orientation (mean±SEM). Gray and black lines/dots in (B-C, H) reflect either individual or grouped measurements (n=4 animals), while asterisks denote significance between groups (*:p<0.05,**:p<0.01, ns: p>0.05).

**Supplemental Figure 6:**
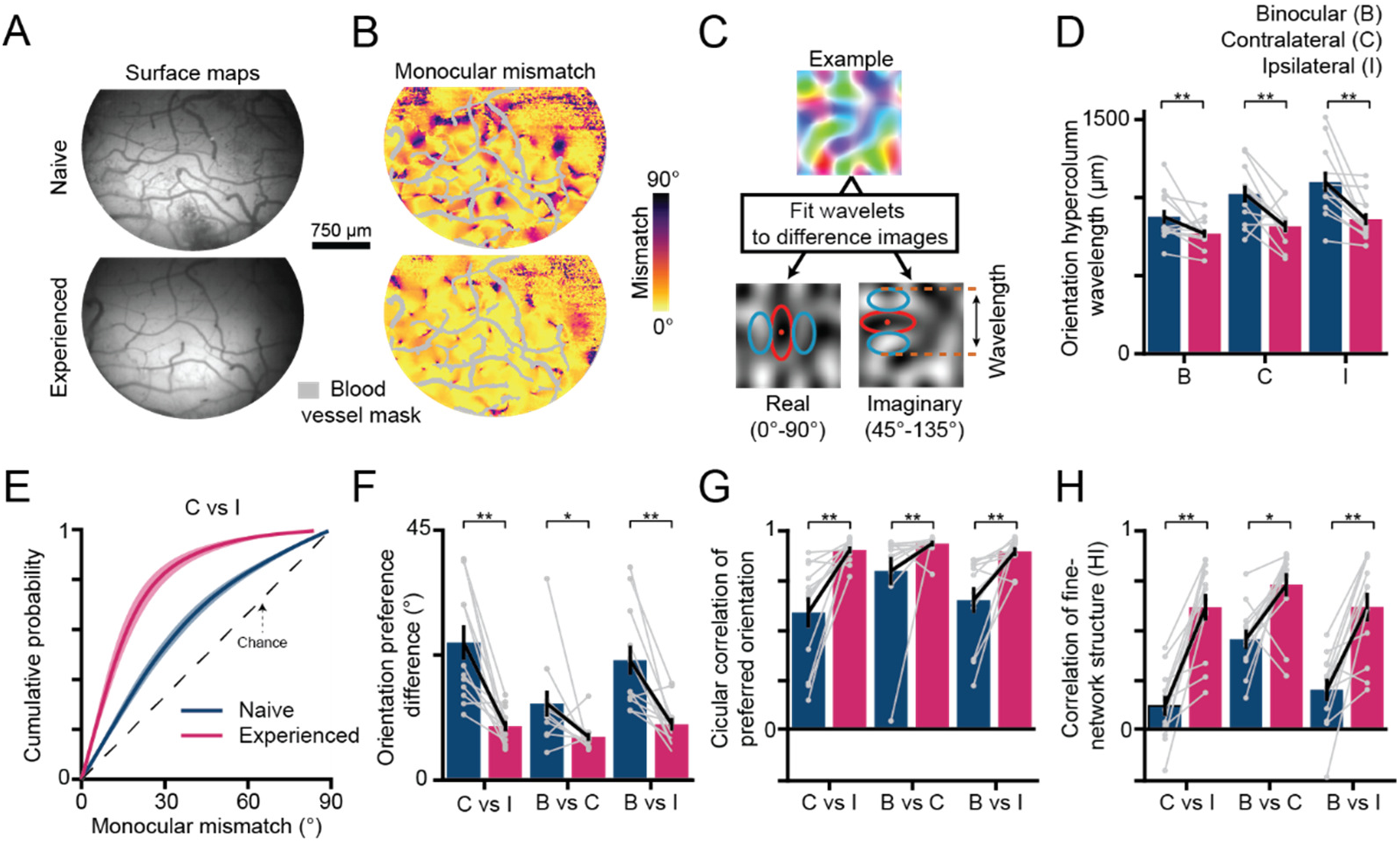
Additional quantification of chronic widefield data. (A) The chronically tracked region before and after eye-opening for the case in Figure 6. (B) The monocular mismatch map for (A). Monocular mismatch is represented by image color, while grayed out regions represent blood vessels. (C) Schematic of fitting wavelets to response maps to measure orientation hypercolumn size. (D) Mean orientation hypercolumn size for chronically tracked monocular and binocular orientation preference maps before and after eye-opening (mean±SEM). Gray and black dots respectively reflect individual or grouped measurements (n=12 animals), while asterisks denote significance differences across time (*:p<0.05,**:p<0.01, ns: p>0.05). (E) Cumulative distributions of monocular mismatch across time for chronically tracked animals (mean±SEM). (F-H) Similar to (D), but comparing the orientation preference difference (F), circular correlation of the preferred orientation (G), and Pearson’s correlation of fine-network structure (H) of the binocular and monocular orientation maps.

## Supplemental Methods

### Viral injections

Viral injections were performed as previously described (Smith et al., 2018). Briefly we expressed GCaMP6s by microinjection of AAV2/1-hSyn-GCaMP6s-WPRE-SV40 (University of Pennsylvania Vector Core) into the visual cortex 6-14 days before imaging experiments. Anesthesia induction was performed using either ketamine (50mg/kg, IM) and/or isoflurane (1-3%) delivered in O_2_ and then maintained with isofluorane (1-2%). Atropine (0.2mg/kg, IM) was administered to reduce secretions, while Buprenorphine (0.01mg/kg, IM) and a 1:1 mixture of lidocaine and bupivacaine (injected directly into the scalp) were administered as analgesics. Animal temperatures were maintained at 37°C using a homeothermic heating blanket. Animals were also mechanically ventilated and both heart rate and end-tidal CO2 were monitored throughout the surgery. Under aseptic surgical technique, a small craniotomy was made over visual cortex 6.5-7 mm lateral and 2mm anterior to lambda. Approximately 1μL of virus was pressure infused into the cortex through a pulled glass pipette across two depths (∼200μm and 400μm below the surface). This procedure reliably produced robust and widespread GCaMP6s expression in excitatory neurons over an area >3 mm in diameter (Smith et al., 2018). To improve the uniformity of the GCaMP6s expression, sometimes an additional injection of 1μL of virus was pressure injected into a separate region of cortex displaced ∼1-1.5mm away from the first injection site.

### Eyelid Suture Procedure

In deprivation and chronic imaging experiments, eyelids were binocularly sutured during a short surgical procedure between P26-P30. Anesthesia was induced with isoflurane (2-5%) and Buprenorphine (0.01mg/kg, IM) was administered. Using aseptic technique, both eyelids were sutured shut using continuous sutures (6-0 Ethilon suture). During recovery, antibiotic ointment was applied to the suture eyelid margins. Eyelid sutures were monitored daily until removed during an imaging experiment as noted in the main text.

### Cranial window surgery

Animals were anesthetized and prepared for surgery as described above. In acute imaging experiments, skin and muscle overlying visual cortex were reflected and a custom-designed metal headplate (8mm DIA) was implanted over the injected region with MetaBond (Parkell Inc.). Then a craniotomy (∼5mm) and a subsequent durotomy were performed, and the underlying brain stabilized with a custom-designed titanium metal cannula (4.5mm DIA, 1.5mm height) adhered to a thin 4mm coverslip (Warner Instruments) with optical glue (#71, Norland Products, Inc). The headplate was hermetically sealed with a stainless-steel retaining ring (5/16” internal retaining ring, McMaster-Carr) and glue (VetBond, 3M or Krazy Glue).

To allow repeated access to the same imaging field in chronic imaging experiments, cranial windows were implanted in each animal 2 days prior to the first imaging session. Using aseptic surgical technique, skin and muscle overlying the dorsal brain surface were gently reflected. To reduce the chances of an implant failure for our cranial window, we used a two-piece design. We adhered a metal headpost (7×7mm) to the skull ∼3.5mm anterior of bregma and a separate custom-designed, chamber implant overlying the injected region using MetaBond and black dental acrylic. Like the acute experiments, we performed a similar craniotomy, durotomy, and inserted the same custom-designed titanium metal cannula. At the end of the survival cranial window implant surgery, the metal cannula was sealed with a silicone polymer plug (Kwik-kast, World Precision Instruments) to protect the imaging window between imaging experiments.

### Calcium Imaging Experiments

Widefield epifluoresence imaging of GCaMP6s was achieved with a Zyla 5.5 sCMOS camera (Andor) controlled by μManager (Edelstein et al., 2010). Images were acquired at 15Hz with 4×4 binning to yield 640×540 pixels through a 4x air-immersion objective (Olympus, UPlanFL 4x N/0.13NA). For analysis, images were spatially downsampled by a factor of 2x to yield 320×270 pixels at a spatial resolution of 11.63µm/pixel. Two-photon imaging of GCaMP6s was performed with a B-Scope microscope (ThorLabs) driven by a Mai-Tai DeepSee laser (Spectra Physics) at 940nm. The B-Scope microscope was controlled by ScanImage (Vidreo Technologies) in a resonant-galvo configuration with multi-plane images (512×512 pixels) being sequentially collected across either one or four imaging planes using a piezoelectric actuator for an effective frame rate of 30Hz or 6Hz respectively. Images were acquired at 2x zoom through a 16x water immersion objective (Nikon, LWD 16X W/0.80NA) yielding cellular fields-of-views of ∼0.7mm on each side (1.36µm/pixel).

Imaging experiments fell into two broad categories: acute and survival imaging experiments. Acute imaging experiments began immediately after the cranial window surgery. For survival imaging experiments, where animals had a cranial window implanted days earlier, anesthesia was induced with isoflurane (2-5%) and atropine (0.2mg/kg) was administered. Animals were intubated and ventilated, and an IV catheter was placed in the cephalic vein. The silicon polymer plug overlying the sealed imaging chamber was gently peeled off. Whenever the imaging quality of the chronic cranial window was found to be suboptimal for imaging, the chamber was opened under aseptic conditions, any regrown tissue/neomembrane was removed, and a new coverslip was inserted.

### Visual stimulation

Visual stimuli were delivered on a LCD screen placed approximately 25– 30cm in front of the eyes using PsychoPy (Peirce, 2007). To evoke orientation responses, full-field square gratings at 100% contrast, at 0.015 or 0.06 cycles per degree and drifting at 1 or 4 Hz were presented at 8 or16 directions. In addition, “blank” stimuli of 0% contrast were also presented. Stimuli were randomly interleaved and presented for 3-4s followed by 3-6s of gray screen.

### General Analysis

All data analysis was performed using custom written scripts in either Python, Matlab, or ImageJ. Early pre-processing steps for analysis were common to both two-photon and widefield epifluorescence imaging data. First, to correct for brain movement during imaging, we registered every imaging frame by maximizing phase correlation to a common reference frame. Furthermore, every imaging experiment acquired during a single day were registered into one reference frame. ROI segmentation was performed manually in widefield epifluorescence imaging by manually drawing around cortical regions with robust visually evoked activity, while cellular ROIs were drawn using custom software in ImageJ (Cell Magic Wand)(Wilson et al., 2017). The raw fluorescence trace for each cellular trace was the average fluorescence trace across all pixels within the ROI. The baseline F_0_ for each pixel/cell was obtained by applying a rank-order filter to the raw fluorescence trace (25-30^th^ percentile) with a time window of 60s. The baseline corrected evoked activity was calculated as *(F-F_0_)/F_0_*= ΔF/F_0_. Grating evoked responses were then calculated as the difference of the average of the fluorescence ΔF/F_0_ over the full stimulus period (3-4s) and the average of the fluorescence ΔF/F_0_ over the pre-stimulus period (1.5-2s). Trial-averaged responses were calculated by taking the median across repeated trials to the same stimulus orientation.

### Widefield Epifluorescence Imaging

For analysis of the wide-field epifluorescence imaging data, spatial filtering was necessary to eliminate signal strength variations and measurement noise. To determine an appropriate high-pass filter cutoff that did not fundamentally alter the underlying modular response, we first computed the average size (i.e. wavelength) of orientation hypercolumns for each imaging experiment using an established wavelet technique (Kaschube et al., 2010). First, we computed a spatially unfiltered orientation preference map using the vector-sum approach:

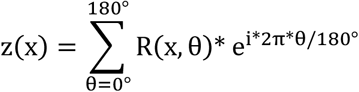

where z(x) is the complex-field orientation response and R(x,θ) is the trial-averaged response to an oriented, drifting grating (θ) at each pixel (x). We then independently fit an array of wavelet coefficients, I(x, θ, l) to the z-scored real- and imaginary-components of z(x):

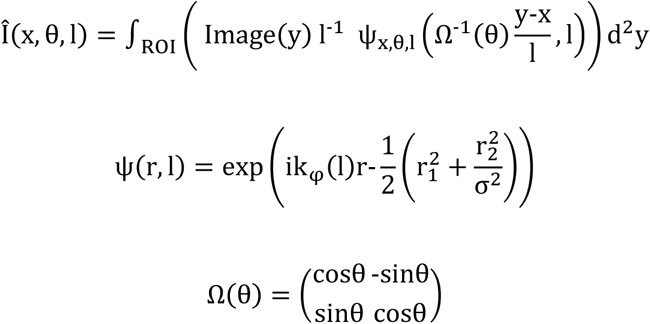

where ψ_x,θ,1_ is the complex Morlet wavelet, Ω(θ) is the 2D rotation matrix, (x, θ, l) are the position, orientation, and scale of the wavelet, and k_φ_ and σ are the characteristic wavenumber and anisotropy of the wavelet. Relatively small, isotropic wavelets (k_φ_ = (2,0) and σ=1) were used to measure the local size or wavelength of orientation hypercolumns . Each wavelet’s orientation (θ) was rotated between 16 uniformly spaced orientations. The wavelet’s scale (l) was varied between a range of [0.4 1.8] mm with a step size of 0.05 mm. The wavelet domain spacing, Λ(x), was computed with the following formulas:

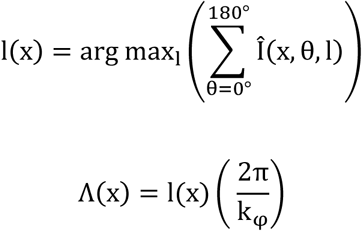

where x is the pixel location and l(x) corresponds to the scale of the wavelet that maximized the angle averaged array of wavelet coefficients (using cubic splines interpolation). The wavelength of orientation hypercolumns, Λ(ROI), for each animal experiment was then defined as the mean wavelet domain spacing over the entire ROI and across both fits to the z-scored real- and imaginary-components of z(x). For quantification of the wavelength in Supplementary Figure 6, the wavelength was computed again, but instead on the vector sum of the Cohen^’^s d (θ, x) spatially filtered, single-condition maps (see below).

We spatially filtered each trial-evoked response image using a spatial bandpass filter, where the low-pass filter cutoff was defined as 50µm and the high-pass filter cutoff was defined as the mean orientation hypercolumn wavelength across monocular and binocular stimulation. A spatial filter was applied to each image in the Fourier Domain, using a filter kernel based on the Fermi-Dirac distribution:

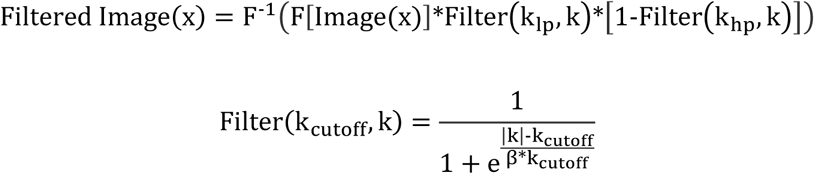

where F denotes the forward Fourier transform, F^-1^ denotes the inverse Fourier transform, k is the spatial wave-number (k = 2π/λ), k_cutoff_ is the filter cutoff (low-pass cutoff (l_lp_): 2π/0.25mm, high-pass cutoff (l_hp_): 2π/1.6mm), and β is the steepness parameter of the fermi-filter (T = 0.05). To maintain the original response amplitude of the population responses, spatially filtered images were ranged-fitted, such that the 5^th^ and 95^th^ percentiles intensity values matched the original, unfiltered images.

To evaluate orientation selective responses, we computed normalized single-condition response maps using the Cohen’s d metric, rather than the customary OSI or 1-CV metrics, to account for response variance across trials:

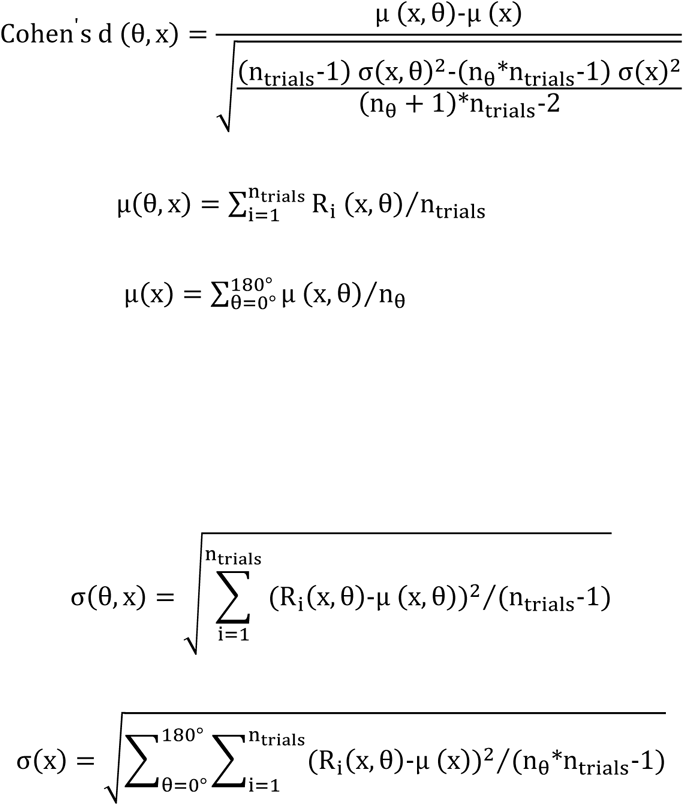

where n_tria1s_ is the number of trials, and n_θ_ is the number of stimulus conditions, and R_i_(x, θ) is the evoked response to an oriented, drifting grating (θ) for trial i at pixel (θ). Preferred orientation (θ_pref(x)_, ranging from 0 to 180°) and orientation selectivity (r) was computed as the vector sum of the Cohen^’^s d (θ, x) single-condition maps:

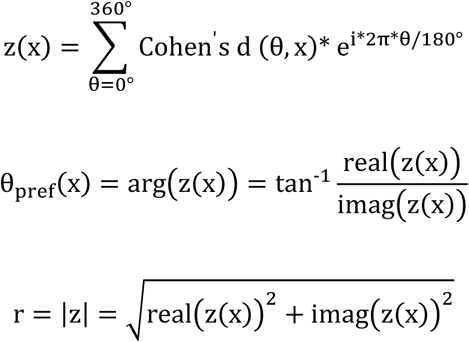

For each pixel, a permutation test (n=100) was used to evaluate if orientation selectivity was significant (p>0.05). Pixels were considered well-tuned if they exhibited statistically significant orientation selectivity for both monocular and binocular stimulation. We excluded any pixels overlying blood-vessels that could be observed in the raw fluorescence images. Monocular mismatch was computed as the absolute angular difference between the contralateral and ipsilateral preferred orientations:

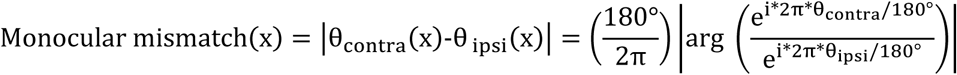

where θ_contra_ and θ_ipsi_ are the preferred orientations for each eye at pixel (x). To ensure that the measured monocular mismatch for the population could not be artifactually explained by high variation in the preferred orientation, we evaluated the worst-case measurement error. Here we bootstrap resampled trial-averaged responses using replacement and recomputed the preferred orientation (n=1,000). The measurement error for each pixel was defined as the median error of the bootstrapped preferred orientations with the measured preferred orientation. We then defined our worst case measurement error as the value of the pixel exhibiting large measurement error (95^th^ percentile).

Orientation preference differences between the monocular and binocular preferred orientations were computed using a similar formula to the monocular mismatch, but switching the preferred orientations of the comparisons. The circular correlation was used to assess differences in the layout of preferred orientation for monocular and binocular orientation maps:

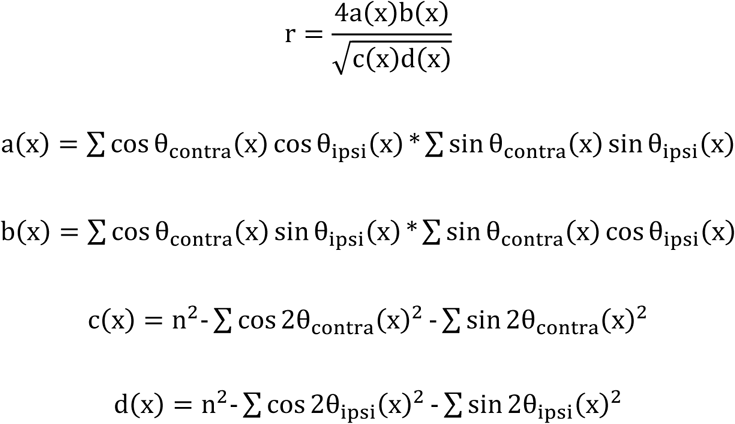

where r is the circular correlation, x is the index of pixels, and n is the number of all orientation selective pixels. For computing the similarity of the fine network structure in orientation preference maps, we computed the Homogeneity Index (Wilson et al., 2017):

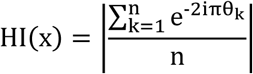

where n is the number of pixels within 100 μm of the seed pixel (x) and θ_k_ is the preference of the k^th^ pixel in the sample. We then quantified the similarity between the Homogeneity Index maps for the monocular and binocular orientation maps using Pearson’s correlation.

### Two-Photon Imaging

Cells were considered responsive if the trial-averaged response to any stimulus orientation for either monocular or binocular stimulation was greater than two standard deviations of the response to a blank stimulus. The preferred orientation for responsive cells was assessed by fitting a Von Mises function to the trial-averaged responses:

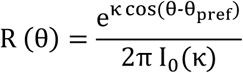

where R (θ) is the trial-averaged response, θ is the stimulation orientation, θ_pref_ is the preferred orientation, κ controls the orientation width, and I_O_(κ) is a zero-order Bessel function. Cells were considered well-tuned if the R^2^ value of the orientation fits to monocular and binocular stimulation were all ≥0.5. To evaluate a cell’s orientation selectivity, we used the Cohen’s d metric and compared the responses of the preferred and orthogonal orientations. Spatial clustering of the preferred orientation or monocular mismatch was assessed by calculating the pairwise correlation for cells located within a fixed range from each other, ranging from distances of 50 µm to 500 µm. To evaluate the degree that all preferred orientations are represented uniformly within each of our cellular fields, we estimated the normalized difference from a uniform distribution using the following metric:

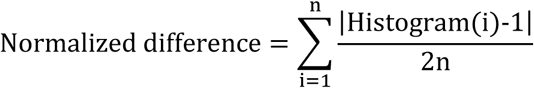

where n is the number of cells, Histogram(i) is the indexed histogram of preferred orientations, ranging from 0° to 180° with a bin size equal to n, i is the index for the histogram bins. The normalized difference metric ranges from 0 (all orientations equally represented) to 1 (only a single representation represented). Monocular mismatch, orientation preference differences, measurement error, and the homogeneity index were computed using the same formulas previously described used for pixels (*see widefield imaging section*).

To evaluate the degree that monocular input from one eye consistently drove stronger responses, we computed the ocular dominance index (ODI):

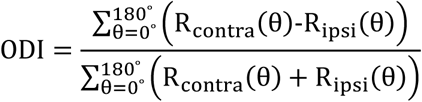

where R_contra_(θ) and R_ipsi_(θ) are the trial-averaged responses to contralateral and ipsilateral stimulation and θ is the stimulation orientation. ODI values range from −1 to +1, where 0 indicates a lack of bias and −1 and +1 values indicate complete dominance from either the ipsilateral or contralateral eyes respectively. Monocularity was defined as the magnitude of the ODI, with values ranging from +0 (binocular) to +1 (monocular).

To decode the stimulus orientation from the population activity vectors recorded during each trial, we used a normalized template matching algorithm (Montijn et al., 2014). In an effort to evaluate the decoding based on the relative responsiveness of each cell, the trial evoked ΔF/F_0_ responses of each cell were first individually z-scored:

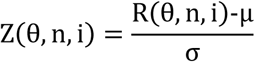

where i indexes the N^th^ neuron, θ is the stimulation orientation, n is the trial number, Z(θ, n, i) is the z-scored response, R(θ, n, i) is the actual ΔF/F_0_ response, µ is the mean ΔF/F_0_ value over the monocular or binocular recording period, and σ is the standard deviation from the mean ΔF/F_0_ value. Template population responses for each stimulation condition (φ) were generated by computing the median across repeated trials to the same stimulus orientation for the z-scored responses. Similarity between the actual trial evoked population vector and each template was defined as:

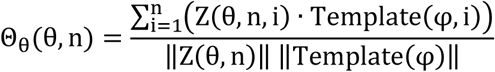

where φ is the stimulus orientation for the template and · indicates multiplication. To ensure our analysis was comparable across different fields-of-view where the number of cells present varies widely, we limited the number of cells to 50 at any time for our decoder. The similarity metric computed for each template was bootstrap resampled without replacement, but a different subset of cells was used for each resampling procedure (n=1,000). The decoded stimulation orientation was defined as the stimulus orientation for the template showing the highest median similarity across bootstrap resampling to the population activity vector.

### Statistics

Unless otherwise noted, hypothesis testing was performed using either a permutation or bootstrapping test. We deliberately chose to use resampling methods for hypothesis testing, rather than relying on standard parametric or non-parametric tests, to avoid confounds that could arise from making assumptions on the underlying sampling distributions present in our data. Statistical significance was assessed by ranking if the observed difference between group averages (i.e. mean or median) was more extreme than the 95^th^ percentile from a surrogate distribution, where the null hypothesis is true. All statistical tests were two-tail, except when assessing if the performance of the normalized template matching decoder was above chance-levels (one-tail). For each hypothesis test, we produced a relevant surrogate distribution by pooling data between groups, bootstrapped new group averages from the pooled data using replacement (n=1,000), and then computed a distribution of differences between surrogate group averages. Unequal group sizes were maintained for unpaired tests, while resampling was restricted between paired measurements for any paired test. Permutation tests were performed similarly, but data was only exchanged between tested groups, rather than resampled with replacement.

## Supplemental statistical note

### Widefield imaging experiments stats

**1. Acute experiments:**

- Naive (n=6 animals, PD27-31): F2053, F2061, F2143, F2145, F2164, F2250
- Experienced (n=7 animals, PD36-42): F2052, F2126, F2127, F2156, F2186, F2249, F2251
**2. Chronic experiments:**

- Chronic (n=12 animals, PD27-38): F2079, F2180, F2182, F2186, F2216, F2217, F2198, F2284, F2303, F2304, F2305, F2317

### Cellular imaging experiments stats

**1. Acute Experiments:**

- Naive (n=3 animals, P28-29): F2143 (n=537 cells), F2145 (n=427 cells), F2164 (n=323 cells)
- Experienced (n=3 animals, PD36-38): F2126 (n=1,272 cells), F2127 (n=423 cells), F2156 (n=439 cells)
- Deprived (n=3, PD35-36): F2144 (n=564 cells), F2150 (n=1,223 cells), F2155 (n=268)
- Recovery (n=5, PD43-46): F2238 (n=1,488), F2245 (n=1,377), F2246 (n=911), F2247 (n=937), F2248 (n=1,217)
**2. Chronic Experiments:**

- Chronic (n=4 animals, PD29-36): F2281 (n=160 tracked cells), F2283 (n=110 tracked cells), F2303 (n=132 tracked cells), F2304 (n=285 tracked cells)

## Figure 1

• **Figure 1f: Pearson’s correlation of monocular, trial-averaged response maps (widefield):** Naive: r=0.31±0.09 vs Experienced: r=0.79±0.03 (Mean±SEM), n=6 vs 7 animals, p<0.001, unpaired permutation test.

• **Figure 1g: Widefield monocular mismatch:** Naive 27.91±4.19° vs Experienced 11.04±1.57° (Mean±SEM), n=6 vs 7 animals, p<0.001, unpaired permutation test.

• **Figure 1j: Cellular monocular mismatch:** Naive 26.87±2.90° vs Experienced 13.11±1.03° (Mean±SEM), n=3 animals, p=0.011, unpaired bootstrap test.

• **Figure 1k: ODI**: Naive 0.05±0.03 vs Experienced 0.23±0.06 (Mean±SD), n=3 animals, p=.04, unpaired bootstrap test.

• **Figure 1n: Spatial clustering by preferred orientation:** Permutation test p-values against random:

| Distance (μm) | [0-50] | 50-100 | 100-150 | 150-200 | 200-250 | 250-300 | 300-350 | 351-400 | 400-450 | 450-500 |
| --- | --- | --- | --- | --- | --- | --- | --- | --- | --- | --- |
| Naive (Contralateral) | <0.001 | <0.001 | 0.998 | >0.999 | >0.999 | >0.999 | 0.998 | 0.996 | 0.878 | 0.990 |
| Experienced (Contralateral) | <0.001 | <0.001 | >0.999 | .998 | .998 | .998 | .998 | .998 | .998 | .998 |
| Naive (Ipsilateral) | <0.001 | <0.001 | 0.998 | >0.999 | >0.999 | >0.999 | >0.999 | >0.999 | 0.998 | >0.999 |
| Experienced (Ipsilateral) | <0.001 | <0.001 | <0.001 | >0.999 | >0.999 | >0.999 | >0.999 | >0.999 | >0.999 | >0.999 |

• **Text**: **Widefield optimal monocular mismatch:** Naive 25.19±4.55° vs Experienced 9.23±1.03° (Mean±SEM), n=6 vs 7 animals, p<0.001, unpaired permutation test.

• **Text: Fraction of responsive cells:** Naive 100±0% vs Experienced 96.27±1.57% (Mean±SEM), n=3 animals.

• **Text**: **Monocularity**: Naive 0.09±0.04 vs Experienced 0.47±0.09 (Mean±SD), n=3 animals, p=.010, unpaired bootstrap test.

• **Text: Fraction of binocularly well-tuned cells:** Naive 56.76±3.94% vs Experienced 71.32±7.75% (Mean±SEM), n=3 animals. Cells well-fit by a Von-Mises fit (R^2^>0.5).

• **Text: Fraction of binocularly well-tuned pixels:** Naive 35.54±4.64% vs Experienced 80.87±2.33% (Mean±SEM), n=6 vs 7 animals.

• **Text: Bootstrapped measurement error: 95^th^ percentile of within condition bootstraps:**

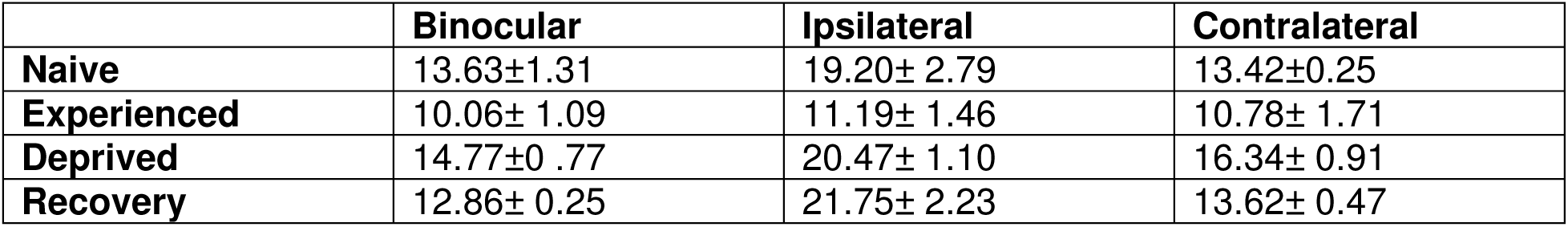

|  | Binocular | Ipsilateral | Contralateral |
| --- | --- | --- | --- |
| Naive | $13.63 \pm 1.31$ | $19.20 \pm 2.79$ | $13.42 \pm 0.25$ |
| Experienced | $10.06 \pm 1.09$ | $11.19 \pm 1.46$ | $10.78 \pm 1.71$ |
| Deprived | $14.77 \pm 0.77$ | $20.47 \pm 1.10$ | $16.34 \pm 0.91$ |
| Recovery | $12.86 \pm 0.25$ | $21.75 \pm 2.23$ | $13.62 \pm 0.47$ |

• **Text: Bootstrapped measurement error: 95^th^ percentile of within condition bootstraps:** Naive θ_contra_=9.46±1.20° and θ_ipsi_=11.01±1.00°, Experienced θ_contra_=8.09±0.54° and θ_ipsi_=7.24±0.44° (Mean±SEM), n=6 or 7 animals.

## Figure 2

• **Figure 2d: Circular correlation of monocular preferred orientations (widefield):** Naive: r=0.44±0.11 vs Experienced: r=0.88±0.03 (Mean±SEM), n=6 vs 7 animals, p<0.001, unpaired permutation test.

• **Figure 2e: Pearson’s correlation of the monocular homogeneity index maps (widefield):** Naive: r=0.41±0.11 vs Experienced: r=0.74±0.03 (Mean±SEM), n=6 vs 7 animals, p<0.001, unpaired permutation test.

• **Figure 2f: Spatial clustering of the monocular mismatch:** Permutation test p-values against random:

| Distance ($\mu\text{m}$) | [0-50] | 50-100 | 100-150 | 150-200 | 200-250 | 250-300 | 300-350 | 350-400 | 400-450 | 450-500 |
| --- | --- | --- | --- | --- | --- | --- | --- | --- | --- | --- |
| Naive | < 0.001 | <0.001 | .0840 | 0.175 | 0.164 | 0.359 | 0.493 | 0.499 | 0.436 | 0.3820 |
| Experienced | <0.001 | 0.5010 | 0.5690 | 0.9040 | 0.7440 | 0.5270 | 0.5700 | 0.5030 | 0.3790 | 0.3090 |

## Figure 3

• **Figure 3d: Cellular monocular mismatch (Deprived vs Experienced):** Deprived 37.75±2.43° vs Experienced 13.11±1.03° (Mean±SEM), n=3 vs 5 animals, p=0.012, unpaired bootstrap test

• **Figure 3d: Cellular monocular mismatch (Recovery vs Deprived):** Recovery 27.00±1.43° vs Deprived 37.75±2.43° (Mean±SEM), n=3 vs 5 animals, p=.016, unpaired bootstrap test

• **Figure 3d: Cellular monocular mismatch (Recovery vs Experienced):** Recovery 27.00±1.43° vs Experienced 13.11±1.03° (Mean±SEM), n=5 vs 3 animals, p=0.012, unpaired bootstrap test

• **Figure 3e: Cellular Cohen’s D (Deprived vs Experienced):** Deprived contralateral 1.03±0.18 vs Experienced contralateral 1.76±0.16 (Mean±SEM), n=3 vs 3 animals, p=0.040, unpaired bootstrap test; Deprived ipsilateral 0.77±0.09 vs Experienced ipsilateral 1.63±0.11 (Mean±SEM), n=3 vs 3 animals, p=0.006, unpaired bootstrap test

• **Figure 3e: Cellular Cohen’s D (Recovery vs Deprived):** Recovery contralateral 0.89±0.04° vs Deprived contralateral 1.03±0.18 (Mean±SEM), n=5 vs 3 animals, p=0.296, unpaired bootstrap test; Recovery ipsilateral 0.75±0.06° vs Deprived ipsilateral 0.77±0.09 (Mean±SEM), n=5 vs 3 animals, p=.702, unpaired bootstrap test

• **Figure 3e: Cellular Cohen’s D (Recovery vs Experienced):** Recovery contralateral 0.89±0.04° vs Experienced contralateral 1.76±0.16 (Mean±SEM), n=5 vs 3 animals, p=0.008, unpaired bootstrap test; Recovery ipsilateral 0.75±0.06° vs Experienced ipsilateral 1.63±0.11 (Mean±SEM), n=5 vs 3 animals, p<0.001, unpaired bootstrap test

• **Figure 3f: Spatial clustering by preferred orientation:** Unpaired bootstrap test versus Naive group:

| Distance (μm) | [0-50) | 50-100 | 100-150 | 150-200 | 200-250 | 250-300 | 300-350 | 350-400 | 400-450 | 450-500 |
| --- | --- | --- | --- | --- | --- | --- | --- | --- | --- | --- |
| Deprived (Contralateral) | .032 | .084 | .464 | .888 | .732 | .942 | .204 | .242 | .420 | 0.888 |
| Deprived (Ipsilateral) | .024 | .028 | .022 | 0.830 | 0.538 | 0.716 | 0.676 | 0.962 | .294 | .400 |
| Recovery (Contralateral) | .004 | .028 | .292 | .964 | 0.642 | 0.820 | 0.07 | 0.086 | 0.252 | 0.770 |
| Recovery (Ipsilateral) | .002 | .006 | .012 | 1 | 0.550 | .5640 | .540 | .914 | .274 | .464 |

• **Figure 3g: Spatial clustering of monocular mismatch:** Unpaired bootstrap test versus Naive group:

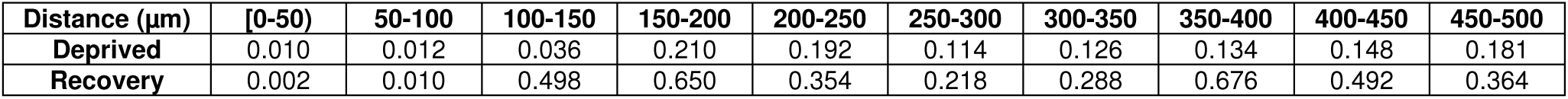

## Figure 4

• **Figure 4h: Cellular orientation preference difference (Binocular vs Contralateral):** Naive 19.12±6.83° vs Experienced 10.44±1.10° (Mean±SEM), n=3 animals, p=.104, unpaired bootstrap test.

• **Figure 4i: Cellular orientation preference difference (Binocular vs Ipsilateral):** Naive 23.40±4.29° vs Experienced 12.73±0.40° (Mean±SEM), n= n=3 animals, p=0.052, unpaired bootstrap test.

• **Figure 4j: Circular correlation of binocular to monocular preferred orientations (widefield):** Naive binocular vs contralateral: r=0.79±0.05 vs Naive binocular vs ipsilateral: r=0.59±0.07 (Mean±SEM), n=6 vs 7 animals, p=0.069, paired permutation test; Experienced binocular vs contralateral: r=0.93±0.01 vs Naive binocular vs ipsilateral: r=0.93±0.02 (Mean±SEM), n=6 vs 7 animals, p=0.91, paired permutation test; Comparison of Naive vs Experienced: Binocular vs contralateral, p=0.018, unpaired permutation test, Binocular vs ipsilateral, p=0.003, unpaired permutation test.

• **Figure 4k: Pearson’s correlation of binocular to monocular homogeneity index (widefield):** Naive binocular vs contralateral: r=0.71±0.03 vs Naive binocular vs ipsilateral: r=0.46±0.12 (Mean±SEM), n=6 vs 7 animals, p=0.035, paired permutation test; Experienced binocular vs contralateral: r=0.82±0.03 vs Naive binocular vs ipsilateral: r=0.82±0.04 (Mean±SEM), n=6 vs 7 animals, p=0.90, paired permutation test. Comparison of Naive vs Experienced: Binocular vs contralateral, p=0.037, unpaired permutation test, Binocular vs ipsilateral, p=0.003, unpaired permutation test.

• **Figure 4m: Fraction of correctly decoded trials (Naive vs Experienced):** Permutation test (see 95^th^ percentile of randomized shuffles)

| Decoding | Actual mean | Shuffle mean | Shuffle 5<sup>th</sup> percentile | Shuffle 95<sup>th</sup> percentile |
| --- | --- | --- | --- | --- |
| Naive binocular | 42.9±5.8 | 12.5 | 8.8 | 16.5% |
| Naive Contralateral | 49.5±5.0 | 12.5 | 8.2 | 17.2% |
| Naive Ipsilateral | 40.2±2.7 | 12.4 | 8.4 | 16.8% |
| Experienced Binocular | 89.1±1.4 | 12.4 | 8.0 | 16.9 |
| Experienced Contralateral | 85.9±2.2 | 12.5 | 8.3 | 17.1 |
| Experience Ipsilateral | 80.1±4.5 | 12.5 | 8.3 | 17.1 |

• **Text and Supplementary Figure 3a: Spatial clustering by preferred orientation for binocular representation:** Permutation test p-values against random or Naive

| Distance (μm) | [0-50) | 50-100 | 100-150 | 150-200 | 200-250 | 250-300 | 300-350 | 350-400 | 400-450 | 450-500 |
| --- | --- | --- | --- | --- | --- | --- | --- | --- | --- | --- |
| Naive (Binocular) | <0.001 | <0.001 | .998 | >0.999 | >0.999 | >0.999 | >0.999 | >0.999 | 0.994 | 0.924 |
| Experienced (Binocular) | <0.001 | <0.001 | >0.999 | .998 | .998 | .998 | .998 | .998 | .998 | .998 |

• **Text and Supplementary Figure 3b: Cellular Cohen’s D (Naive vs Experienced):** Naive binocular 1.13±0.13, contralateral 1.10±0.15, ipsilateral 0.78±0.04, binocular vs contralateral: p = 0.794, binocular vs ipsilateral: p=0.994, contralateral vs ipsilateral: p=0.937; Experienced binocular 1.98±0.07, contralateral 1.76±0.13, ipsilateral 1.63±0.09 (Mean±SEM), binocular vs contralateral p =0.993, binocular vs ipsilateral: p=0.993, contralateral vs ipsilateral: p=0.937. n=3 vs 3 animals.

• **Text and Supplementary Figure 3c: Pearson’s correlation of cellular monocular mismatch vs Binocular Cohen’s D:** Naive r=-0.25±0.12 vs Experienced r=-0.21±0.02 (Mean±SEM), n=3 animals.

• **Text and Supplementary Figure 3d: Pearson’s correlation of cellular orientation preference difference vs ODI:** Binocular vs Contralateral r=-0.20, Binocular vs Ipsilateral r=-0.20.

## Figure 5

• **Figure 5e: Cellular monocular mismatch:** Naïve 35.47±6.09° vs Experienced 12.73±0.94° (Mean±SEM), n=4 animals, p=0.007, paired bootstrap test.

• **Figure 5g: Cellular monocularity (Well-fit cells across binocular and monocular stimulation for both experiment days)**: Naive 0.136±0.009 vs Experienced 0.154±0.003 (Mean±SEM), n=4 animals, p=0.03, paired bootstrap test.

• **Text: Cellular monocularity (Well-fit cells across binocular and monocular stimulation at eye-opening, but not after eye-opening)**: Naive 0.140±0.033 vs Experienced 0.247±0.09 (Mean±SEM), n=4 animals, p=0.38, paired bootstrap test.

• **Text: Pearson’s correlation between Naive monocular mismatch and Experienced monocularity:** r_actual_=0.03 [-0.12 0.17], median [CI], vs r_shuffle_ = 0.14 [-0.02 0.30], bootstrapped correlation.

• **Text and Supplementary Figure 3c: Orientation preference difference between binocular and contralateral:** Naïve 31.06±5.92° vs Experienced 11.25±1.37° (Mean±SEM), n=4 animals, p=0.008, paired bootstrap test.

• **Text and Supplementary Figure 3F: Orientation preference difference between binocular and contralateral:** Naïve 22.01±5.10° vs Experienced 10.17±1.45° (Mean±SEM), n=4 animals, p=0.007, paired bootstrap test.

• **Text and Supplementary Figure 4b: Fraction of correctly decoded trials using Naive Decoder:** Naive binocular 40.8±3.5%, contralateral 39.1±2.0%, ipsilateral 42.3±2.3%, Experienced binocular 24.9±2.9%, contralateral 20.4±3.0%, ipsilateral22.79±2.8% (Mean±SEM), Shuffles: 12.5% [8.13, 16.88] (Mean [CI]). Naive: binocular vs contralateral p=0.620, binocular vs ipsilateral p=0.345, contralateral vs ipsilateral p=0.313, Experienced: binocular vs contralateral p=0.168, binocular vs ipsilateral p=0.007, contralateral vs ipsilateral p=0.402, paired bootstrap test.

• **Text and Supplementary Figure 4c: Fraction of correctly decoded trials using Experienced Decoder:** Naive binocular 21.5±3.1%, contralateral 17.92±2.0%, ipsilateral20.82±2.4%, Experienced binocular 73.4±0.7%, contralateral 66.4±3.7%, ipsilateral 68.1±1.4% (Mean±SEM), Shuffles: 12.5% [8.13, 16.88] (Mean [CI]). Naive: binocular vs contralateral p=0.287, binocular vs ipsilateral p=0.393, contralateral vs ipsilateral p=0.265, Experienced: binocular vs contralateral p=0.030, binocular vs ipsilateral p=0.007, contralateral vs ipsilateral p=0.573, paired bootstrap test.

• **Text and Supplementary Figure 4c-d: Comparison of decoding performance between naive and experienced decoders:** Naive binocular: p=0.007, contralateral: p=0.007, ipsilateral: p=0.007. Experienced binocular: p=0.007, contralateral: p=0.007, ipsilateral: p=0.007.

• **Text and Supplementary Figure 4g: Average shifts in preferred orientation between Naive and Experienced:** binocular 31.14±4.12°, contralateral 30.06±4.06°, ipsilateral 30.13±4.78° (Mean±SEM), n=4 animals. Binocular vs contralateral p=0.84, binocular vs ipsilateral p=0.55, contralateral vs ipsilateral p=0.94, paired bootstrap test.

• **Text and Supplementary Figure 4g: Normalized difference from a uniform distribution:** Naive binocular 0.51±0.05, contralateral 0.65±0.02, ipsilateral 0.66±0.03. Binocular vs contralateral p=0.026, binocular vs ipsilateral p=0.006, contralateral vs ipsilateral p=0.472. Experienced binocular 0.46±0.01, contralateral 0.46±0.02, ipsilateral0.47±0.03. Binocular vs contralateral p=0.678, binocular vs ipsilateral p=0.542, contralateral vs ipsilateral p=0.382, paired bootstrap test.

## Figure 6

• **Figure 6d: Circular correlation of preferred orientation (Naive vs Experienced, chronic widefield):** binocular r=0.40±0.04, contralateral: r=0.30±0.04, ipsilateral r=0.16±0.04 (Mean±SEM), n=12 animals); binocular vs contralateral p=0.024, binocular vs ipsilateral p=0.001, contralateral vs ipsilateral p=0.058, paired bootstrap test.

• **Figure 6e: Pearson’s correlation of homogeneity index (Naive vs Experienced, chronic widefield):** binocular r=0.36±0.05, contralateral: r=0.22±0.05, ipsilateral r=0.09±0.04 (Mean±SEM), n=12 animals); binocular vs contralateral p=0.010, binocular vs ipsilateral p=0.003, contralateral vs ipsilateral p=0.098, paired bootstrap test.

• **Figure 6g: Circular correlation of preferred orientation (Naive binocular/monocular versus Experienced monocular, chronic widefield):** Naive contralateral vs Experienced contralateral (C_N_-C_E_): r=0.30±0.04, Naive binocular vs Experienced contralateral (B_N_-C_E_): r=0.38±0.05, Naive ipsilateral vs Experienced ipsilateral (I_N_-I_E_): r=0.16±0.04, Naive binocular vs Experienced ipsilateral (B_N_-I_E_): r=0.34±0.04 (Mean±SEM), n=12 animals); C_N_-C_E_ vs B_N_-C_E_, p=0.048, I_N_-I_E_ vs B_N_-I_E_, p=0.004, paired bootstrap test.

• **Figure 6h: Pearson’s correlation of homogeneity index (Naive binocular/monocular versus Experienced monocular, chronic widefield):** Naive contralateral vs Experienced contralateral (C_N_-C_E_): r=0.22±0.05, Naive binocular vs Experienced contralateral (B_N_-C_E_): r=0.34±0.05, Naive ipsilateral vs Experienced ipsilateral (I_N_-I_E_): r=0.09±0.04, Naive binocular vs Experienced ipsilateral (B_N_-I_E_): r=0.27±0.04 (Mean±SEM), n=12 animals); C_N_-C_E_ vs B_N_-C_E_, p=0.020, I_N_-I_E_ vs B_N_-I_E_, p=0.018, paired bootstrap test.

• **Text and Supplementary Figure 6d: Mean orientation hypercolumn wavelength**: Naive: λ_binoc_=876.2±45.4μm, λ_contra_=1021.7±54.2μm, λ_ipsi_=1097.9±65.1μm; binocular vs contralateral: p=0.001, binocular vs ipsilateral: p=0.019, contralateral vs ipsilateral: p=0.37; Experienced: λ_binoc_=767.9±27.2μm, λ_contra_=815.2±37.4μm, λ_ipsi_=861.4±39.3μm; binocular vs contralateral: p=0.010, binocular vs ipsilateral: p=0.019, contralateral vs ipsilateral: p=0.33. Naive vs Experienced comparisons: binocular: p=0.001, contralateral: p=0.001, ipsilateral: p=0.001, n=12 animals, paired bootstrap test.

• **Supplementary Figure 6f: Mean orientation preference difference:** Naive: contralateral vs ipsilateral difference 22.25±2.74°, binocular vs contralateral difference 13.72±2.71°, binocular vs ipsilateral 20.11±2.50°; Experienced: contralateral vs ipsilateral difference 9.30±0.93°, binocular vs contralateral difference 7.31±0.73°, binocular vs ipsilateral 9.66±1.14°. Naive vs Experienced comparisons: contralateral vs ipsilateral: p=0.001, binocular vs contralateral: p=0.01, binocular vs ipsilateral: p=0.001, n=12 animals, paired bootstrap test.

• **Supplementary Figure 6g: Circular correlation of the preferred orientation:** Naive: contralateral vs ipsilateral difference r=0.59±0.08, binocular vs contralateral difference r=0.78±0.07°, binocular vs ipsilateral r=0.65±0.07; Experienced: contralateral vs ipsilateral difference r=0.90±0.02, binocular vs contralateral difference r=0.94±0.02, binocular vs ipsilateral r=0.90±0.02. Naive vs Experienced comparisons: contralateral vs ipsilateral: p=0.001, binocular vs contralateral: p=0.003, binocular vs ipsilateral: p=0.001, n=12 animals, paired bootstrap test.

• **Supplementary Figure 6g: Pearson’s correlation of HI:** Naive: contralateral vs ipsilateral difference r=0.12±0.05, binocular vs contralateral difference r=0.46±0.05°, binocular vs ipsilateral r=0.20±0.06; Experienced: contralateral vs ipsilateral difference r=0.63±0.07, binocular vs contralateral difference r=0.74±0.06, binocular vs ipsilateral r=0.63±0.07. Naive vs Experienced comparisons: contralateral vs ipsilateral: p=0.001, binocular vs contralateral: p=0.014, binocular vs ipsilateral: p=0.002, n=12 animals, paired bootstrap test.

